# Biosynthesis of cannabinoid precursor olivetolic acid by overcoming rate-limiting steps in genetically engineered *Yarrowia lipolytica*

**DOI:** 10.1101/2021.06.10.447928

**Authors:** Jingbo Ma, Yang Gu, Peng Xu

**Affiliations:** Department of Chemical, Biochemical and Environmental Engineering, University of Maryland, Baltimore County, Baltimore, MD 21250; School of Food Science and Pharmaceutical Engineering, Nanjing Normal University, Nanjing, China; Department of Chemical Engineering, Guangdong Technion-Israel Institute of Technology, Shantou, Guangdong 515063

**Keywords:** Metabolic Engineering, Oleaginous yeast, Cannabinoids, Olivetolic acid, Natural Products, Biosynthesis

## Abstract

Natural products acting on our central nervous systems are in utmost demand to fight against pain and mental disorders. Cannabinoids (CBDs) are proven neuroactive agents to treat anxiety, depression, chronic pain diseases, seizure, strokes and neurological disorders. The scarcity of the hemp-sourced CBD products and the prohibitive manufacturing cost limit the wide application of CBDs. Yeast metabolic engineering offers the flexibility to meet the ever-increasing market demand. In this work, we took a retrosynthetic approach and sequentially identified the rate-limiting steps to improve the biosynthesis of the CBD precursor olivetolic acid (OLA) in *Yarrowia lipolytica*. We debottlenecked the critical enzymatic steps to overcome the supply of hexanoyl-CoA, malonyl-CoA, acetyl-CoA, NADPH and ATPs to redirect carbon flux toward OLA. Implementation of these strategies led to an 83-fold increase in OLA titer in shaking flask experiment. This work may serve as a baseline for engineering CBD biosynthesis in oleaginous yeast species.

## 1. Introduction

Cannabinoids (CBDs) are a large family of neurological active compounds. Cell biologists have identified multiple cannabinoids receptor proteins in human brain and found CBDs or their derivatives played an important role to regulate human cognitive and emotional functions ^1, 2^. Recent pharmaceutical and clinical studies have proved that CBDs can be used to treat anxiety, depression, aging-related muscle/joint pain, seizure, stroke and cardiovascular diseases *et al* ^3, 4^. Almost all CBDs are sourced from farm-harvested hemp or *Cannabis sativa*-related species. Recent molecular breeding has resulted in special hemp cultivars which synthesize less tetrahydrocannabinol (THC) -- the principle psychoactive constituent of cannabis (*Marijuana*) ^5^. CBD oils free of THCs will not cause human addiction. FDA and World Health Organization (WHO) have legally approved the use of CBDs as an essential drug ingredient to treat neurological disorders. Recently, a growing number of nations have lifted the restrictions and approved the use of THC-free CBDs as a consumer chemical in human food, drink, cosmetics and nutraceutical industry. For example, hemp-derived CBD oil (free of THC) has been infused with human lotion, confectionary (gummies or cookies) or cigarette products. The overall market size of CBD oil as consumer chemicals is estimated to be $25 billion in 2025 (assuming 50 Million people will use CBDs as a consumer chemical daily, average dosage 50g/people/year or 2 mg/kg-body-weight/day). Current crude CBD oil (45% strength) is sold at about $3,000/kg and pure CBD oil is sold at $0.01 per milligram. CBD market is projected to increase 10%-15% annually, the current hemp planting and agricultural technology cannot meet this rapid market demand. The pandemics of Covid-19 has severely disrupted our supply chain of food, pharmaceuticals and consumer chemicals, driving us to seek alternate solutions to manufacture essential drugs and safeguard our well-being ^6^. There is an urgent need to develop alternate route to fill the CBD supply chain.

Microbial metabolic engineering is considered as the enabling technology to mitigate environmental concerns and address the scarcity of resource limitation challenges ^7, 8^. Compared to plant-based production system, microbes have a number of advantages, including less dependence on arable land or climate changes, ease of genetic manipulation and large-scale production, as well as robust growth and high conversion rate across a wide range of low-cost renewable raw materials ^6, 9^. Therefore, genetically modified microbes have been widely used to produce fuels, commodity chemicals and nutraceuticals ^10^. With our increased knowledge of cellular physiology and understanding of molecular genetics toward higher and complex organisms, there is a growing interest to develop novel cellular chassis that may overcome the constraints of common host (*E. coli* and *S. cerevisiae*) ^11^. For example, *Yarrowia lipolytica*, is characterized as a generally regarded as safe (GRAS) oleaginous yeast, has been recently modified to produce an arrange of value-added natural products, including resveratrol ^12^, squalene ^13^, flavonoids ^14^, artemisinin ^15^ and violacein ^16^ *et al*. The distinct oil-accumulating (hydrophobic) environment, abundance of intracellular membrane surface, and the compartmentalization of biosynthetic pathways in *Y. lipolytica* provide the ideal microenvironment for the catalytic functions of enzymes with stereo- or regio-selectivity ^11^. This property is extremely important for the functional expression of P450 enzymes with site-specific hydroxylation or peroxidation reactions. CBD biosynthetic pathway involves polyketide synthase, mevalonate pathway and prenyltransferase, which posts a significant challenge for efficient biosynthesis in common host organism ^17^. *Y. lipolytica* is reported to accommodate strong flux for acetyl-CoA, malonyl-CoA and HMG-CoA. Naturally, *Y. lipolytica* could be a promising host to produce CBDs and their derivatives. In this work, we explored the possibility to use oleaginous yeast *Y. lipoltyica* as the host to synthesize olivetolic acid, a universal precursor to synthesize CBDs. We overcome a number of critical pathway bottlenecks to unlock the potential of *Y. lipolytica* to synthesize olivetolic acids. These strategies, when combined, lead to a more than 83-fold increase in olivetolic acid production. This preliminary result may provide a baseline for us to develop CBD-producing oleaginous yeast cell factories.

## 2. Material and Methods

### 2.1. Strains, plasmids, primers, and chemicals

All strains of engineered *Y. lipolytica*, including the genotypes, recombinant plasmids, and primers have been listed in Supplementary Table 1 and 2. Olivetolic acid was purchased from Santa Cruz Biotechnology. All other chemicals were obtained from Sigma-Aldrich and Fisher Scientific. Heterologous synthetic genes including genes *CsOLS, CsOAC, CsAAE1, CsAAE3, PpLvaE, SeACS*^L641P^ and *McMAE2* were codon-optimized using the online IDT Codon Optimization Tool and then ordered from GENEWIZ (Suzhou, China). The synthetic gene fragments were assembled with the New England Biolab Gibson Assembly kits, with the pYaliBrick vector as plasmid backbone. DNA sequences were verified by Sanger sequencing (QuintaraBio).

### 2.2. Shake flask cultivations and pH control

For performing shake flask cultivations, seed culture was carried out in the shaking tube with 2 mL seed culture medium at 30 °C and 250 r.p.m. for 48 h. Then, 0.6 mL of seed culture was inoculated into the 250 mL flask containing 30 mL of fermentation medium and grown under the conditions of 30 °C and 250 r.p.m. for 96 h. One milliliter of cell suspension was sampled every 24h for OD_600_ and desired metabolite measurement.

Seed culture medium used in this study included the yeast complete synthetic media regular media (CSM, containing glucose 20.0 g/L, yeast nitrogen base without ammonium sulfate 1.7 g/L, ammonium sulfate 5.0 g/L, and CSM-Leu 0.74 g/L) and complex medium (YPD, containing glucose 20.0 g/L, yeast extract 10.0 g/L, and peptone 20.0 g/L). Fermentation medium used in this study contained the yeast complete synthetic media regular media (CSM, containing glucose 40.0 g/L, yeast nitrogen base without ammonium sulfate 1.7 g/L, ammonium sulfate 1.1 g/L, and CSM-Leu 0.74 g/L) and complex medium (YPD, containing glucose 40.0 g/L, yeast extract 10.0 g/L, and peptone 20.0 g/L). To control the pH, 20 mM phosphate buffer saline (PBS, Na_2_HPO_4_-NaH_2_PO_4_) or 20 g/L CaCO_3_ was used, respectively.

### 2.3. Yeast transformation and screening of high-producing strains

The standard protocols of *Y. lipolytica* transformation by the lithium acetate method were described as previously reported ^16, 18^. In brief, one milliliter cells was harvested during the exponential growth phase (16-24 h) from 2 mL YPD medium (yeast extract 10 g/L, peptone 20 g/L, and glucose 20 g/L) in the 14-mL shake tube, and washed twice with 100 mM phosphate buffer (pH 7.0). Freshly cultivated yeast colony lawns picked from overnight-grown YPD plates could also be used for genetic transformation. Then, cells were resuspended in 105 μL transformation solution, containing 90 uL 50% PEG4000, 5 μL lithium acetate (2M), 5 μL boiled single stand DNA (salmon sperm, denatured) and 5 μL DNA products (including 200-500 ng of plasmids, linearized plasmids or DNA fragments), and incubated at 39 °C for 1 h, then spread on selected plates. It should be noted that the transformation mixtures needed to be vortexed for 15 seconds every 15 minutes during the process of 39 °C incubation. The selected markers, including leucine, uracil and hygromycin, were used in this study. All engineered strains after genetic transformation were undergone PCR screening using the GoTaq Green PCR kits, and the strain containing the correct gene fragment was selected to perform shaker flask cultivation. For shaking tube cultivations, 100 μL seed cultures were inoculated into 5 mL fermentation media in a 50 mL tube.

### 2.4. Single-gene and multi-genes expression vectors construction

In this work, the YaliBrick plasmid pYLXP’ was used as the expression vector ^19^. The process of plasmid constructions followed the YaliBrick gene assembly platforms ^16^. In brief, recombinant plasmids of pYLXP’-*xx* (a single gene) were built by Gibson Assembly of linearized pYLXP’ (digested by *SnaBI* and *KpnI*) and the appropriate PCR-amplified or synthetic DNA fragments. Multi-genes expression plasmids were constructed based on restriction enzyme subcloning with the isocaudamers *AvrII* and *NheI*. All genes were respectively expressed by the TEF promoter with intron sequence and XPR2 terminator, and the modified DNA fragments and plasmids were sequence-verified by Sanger sequencing (Quintarabio).

### 2.5. Gene knockout

A marker-free gene knockout method based on Cre-*lox* recombination system was used as previously reported ^16, 20^. For performing gene knockout, the upstream and downstream sequences (both 1000 bp) flanking the deletion targets were PCR-amplified. These two fragments, the *loxP*-*Ura/Hyr*-*loxP* cassette (digested from plasmid pYLXP’-l*oxP-Ura/Hyr* by *AvrII* and *salI*), and the gel-purified plasmid backbone of pYLXP’(linearized by *AvrII* and *salI*) were joined by Gibson Assembly, giving the knockout plasmids pYLXP’-*loxP*-*Ura/Hyr*-*xx* (xx is the deletion target). Next, the knockout plasmids were sequence-verified by Quintarabio. Then, the gene knockout cassettes were PCR-amplified from the knockout plasmids pYLXP’-*loxP*-*Ura/Hyr*-*xx*, and further transformed into *Y. lipolytica*. The positive transformants were determined by colony PCR. Knockout strains were built on top of the Ku70-deficient strains (Po1f background). Subsequently, plasmid pYLXP’-*Cre* was introduced into the positive transformants and promoted the recombination of *loxP* sites, which recycle the selected marker. Finally, the intracellular plasmid pYLXP’-*Cre* was evicted by incubation at 30°C in YPD media for 48h. Here, *Ura* is the uracil marker, and *Hyr* is hygromycin marker.

### 2.6. Genomic integration of desired genes

In this work, genomic integration of desired genes was performed in two different ways: site-specific genomic integration plasmids or application of pBR docking platform by linearizing the plasmid pYLXP’ with digested enzyme *NotI*. Here, we constructed two genomic integration plasmids pURLA and pURLB, corresponding to the *Ku70* and *YALI0C05907g* (encoding a hypothetical protein conserved in the *Yarrowia* clade) genomic sites, respectively. The procedure of using these two plasmids was similar as that of gene knockout protocol. The method of constructing integration plasmids was described in previous work. The application of pBR docking platform was achieved by linearizing the plasmid pYLXP’ with *NotI* restriction enzyme digestion.

### 2.7. HPLC quantification and LC-MS characterization of olivetolic acid

Cell densities were monitored by measuring the optical density at 600 nm (OD_600_). The concentrations of olivetolic acid were measured by high-performance liquid chromatography (HPLC) through Agilent HPLC 1220. In detail, olivetolic acid was measured at 270 nm under 40 °C (column oven temperature) with a mobile phase containing 60% (v/v) methanol in water at a flow rate of 0.4 mL/min equipped with a ZORBAX Eclipse Plus C18 column (4.6 × 100 mm, 3.5 μm, Agilent) and the VWD detector.

To quantify the concentration of olivetolic acid, 0.5 mL whole cell sample with both cell pellet and liquid culture was taken. Subsequently, samples were treated with 2 U/OD zymolyase (2h, 30 °C with shaking at 1000 rpm), and then cell suspensions were added with 20% (w/v) glass beads (0.5 mm) and the cells were grinded with hand-powered electrical motor (VWR). Subsequently, the crude extracts were mixed with an equal volume of ethyl acetate (v/v), followed by vortex at room temperature for 2 hours. Organic and inorganic layers were separated by centrifugation at 12000 rpm for 10 min. Samples were extracted three times. The combined organic layers were evaporated in a vacuum oven (50 °C) and the remainders were resuspended in 0.5 mL 100% methanol. Then, 100 uL of the sample were gently transferred into a HPLC vial insert and 5 uL were injected into HPLC for OLA quantification. Under this condition, the retention time for OLA is 10.8 min.

Olivetolic acids standards and samples were also characterized by the Perkin Elmer QSight LX50 UHPLC (ultra-high performace liquid chromatography) with the QSight 210 Mass Spectrometer under negative ESI (electrospray ionization) mode. The Agilent Zorbax Eclipse XDB-C18 2.1 × 50mm, 1.8 μm (P/N 981757-902) column was used for the UHPLC system. Under the UHPLC-MS system, the retention time for olivetolic acid is 1.8 min and the characteristic mass/charge (m/z) ratio for OLA is 223.5 and 179.5 (decarboxylated-CO_2_ fragment). A detailed UPLC-MS report for OLA could be found in the supplementary information.

## 3. Results and Discussion

### 3.1 Biosynthesis of olivetolic acid in engineered Y. lipolytica

Olivetolic acid (OLA) biosynthesis involves the condensation of one hexanoyl-CoA with three malonyl-CoAs by a type III PKS (tetraketide synthase) OLA synthase (*C. sativa* OLS; *Cs*OLS) and an OLA cyclase (*Cs*OAC) (Figure 1), both enzymes were derived from the *Cannabis* species. Previous studies have achieved trace amount of OLA production in both *E. coli* and *S. cerevisiae* ^21, 22^. To produce OLA in *Y. lipolytica*, the plasmid pYLXP’-*CsOLS*-*CsOAC* encoding the codon-optimized *CsOLS* and *CsOAC* was transformed into *Y. lipolytica*. Under HPLC characterization, OLA has a characteristic retention time of 10.8 min (Figure 2a). The resulting strain YL101 only produced 0.11 mg/L OLA after 96 h cultivation (Figure 2b), which is comparable to the initial OLA production in *S. cerevisiae*.

**Figure 1.**
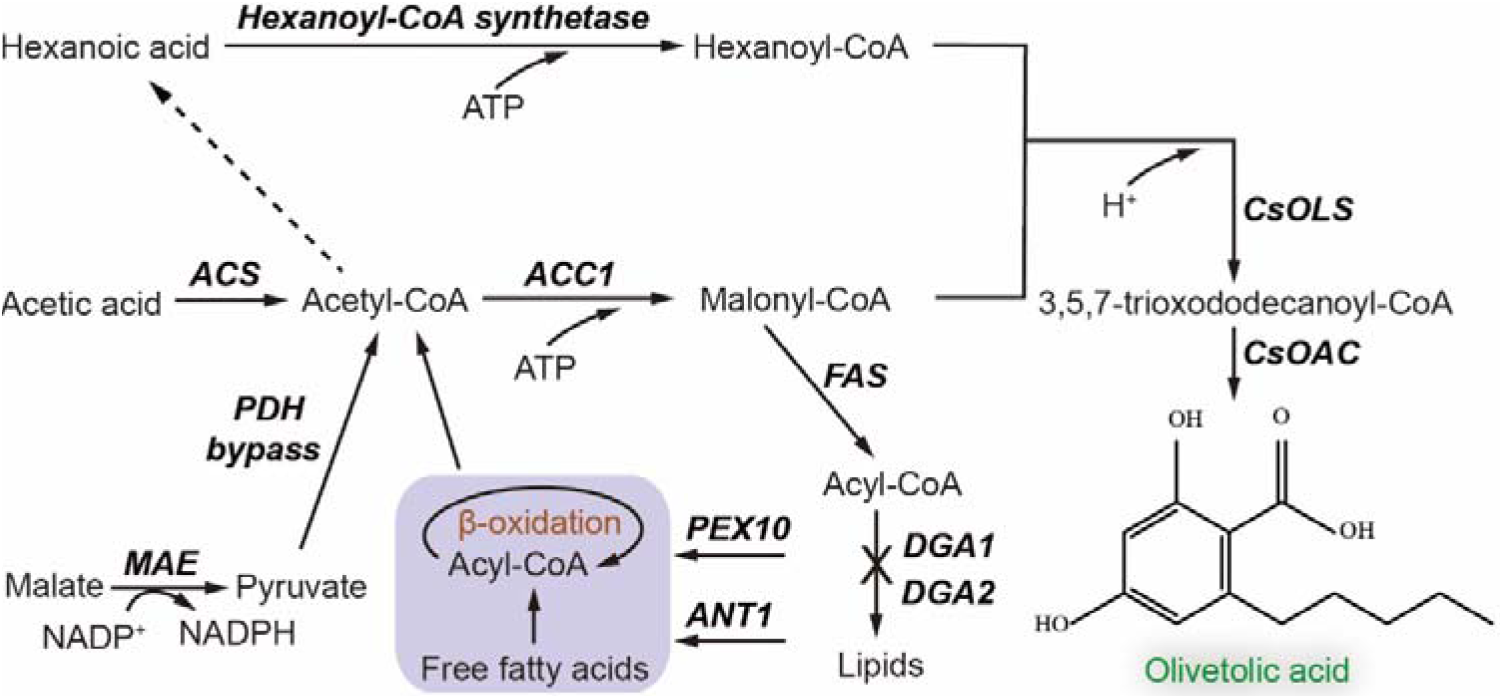
Engineered metabolic pathway for synthesis of olivetolic acid (OLA) in *Y. lipolytica*. ACC1, acetyl-CoA-carboxylase; ACS, acetic acid synthase; ANT1, adenine nucleotide transporter; CsOAC, OLA cyclase from *Cannabis sativa*; CsOLS, OLA synthase from *C. sativa*; DGA1 and DGA2, diacylglycerol acyltransferases; FAS, fatty acid synthase; MAE, malic enzyme; PDH, pyruvate dehydrogenase; PEX10, peroxisomal matrix protein.

**Figure 2.**
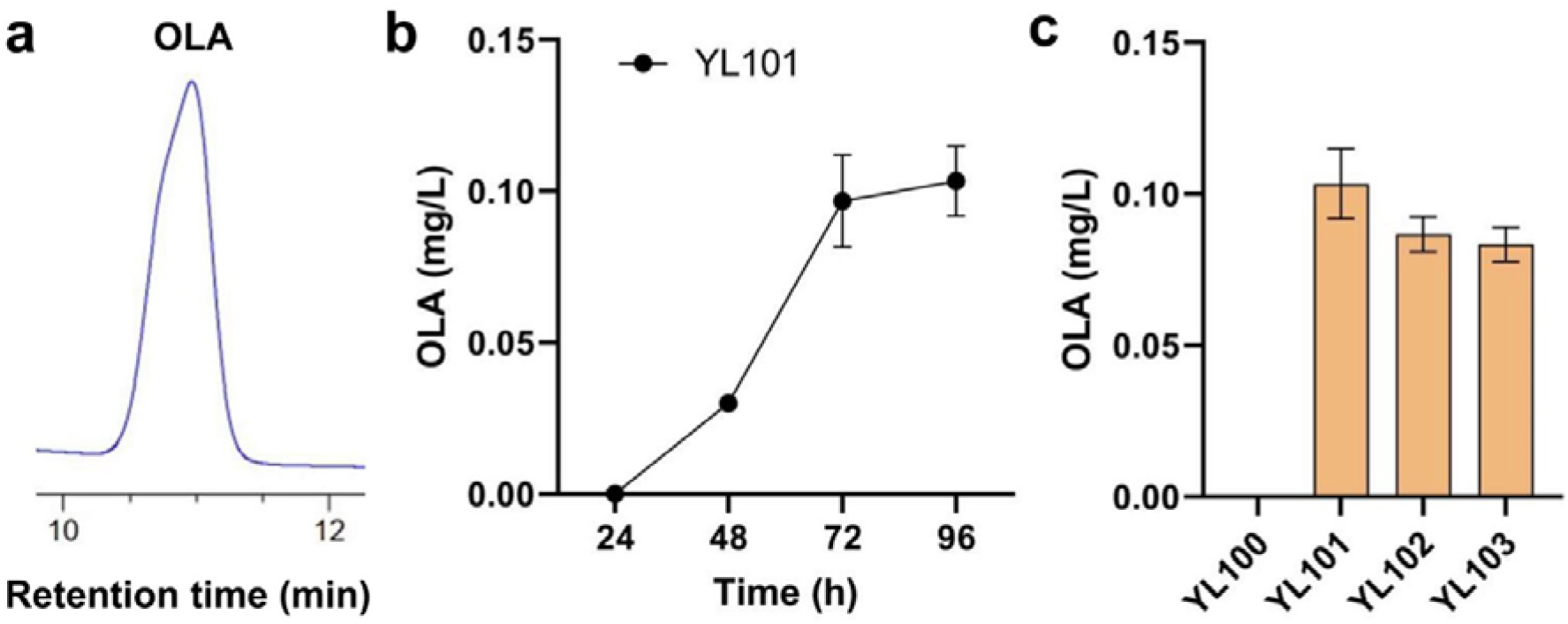
Production of olivetolic acid (OLA) by recruiting OLS and OAC. (a) *In vivo* production of OLA in YL101 strain. (**a**) Extracts were analyzed by HPLC and signals were compared to authentic OLA standards (stds). (**b**) Time profiles of OLA titer in YL101 strain. (**c**) OLA production in engineered strains by expressing OLS and OAC or their fusions.

Evidence showed that some undesired byproducts could also be formed in the biosynthetic pathway of OLA ^21, 22^. We hypothesized that fusing *CsOLS* with *CsOAC* may place the two enzymes in close proximity and minimize the dissipation of intermediates. In such a way, fused OLS-OAC may efficiently transfer the intermediates to form OLA with minimal byproducts. We fused the two proteins with an amino acid linker (10×Glycine) in two different orientations. The results showed that when *CsOLS* was fused to the N-terminus or C-terminus of the *CsOAC*, the obtained strains YL102 and YL103 showed a slightly decline in the production of OLA compared with the control strains YL101 (Figure 2c). It is possible that the fusion of these two proteins may negatively impact their correct folding and catalytic functions. Because the fusion of *CsOLS* with *CsOAC* could not further improve OLA production, the best production strain, YL101, was subjected to further engineering.

### 3.2 Screening enzymes to overcome the limitation of precursor hexanoyl-CoA supply

One of the critical challenges to improve OLA biosynthesis is the pool size of hexanoyl-CoA, which was proven to be a rate-limiting factor inside both *E. coli* and *S. cerevisiae* ^21, 22^. An acyl activating enzyme encoded by *CsAAE1* from *Cannabis sativa* was characterized to catalyze the formation of hexanoyl-CoA from hexanoic acid, and the expression of *Cs*AAE1 has been shown to successfully increase the titer of OLA by twofold in *S. cerevisiae* with the feeding of 1 mM hexanoic acid ^22^. Hexanoic acid is toxic to cell growth and inhibit enzyme activity. The dosage of hexanoic acid should be investigated to minimize its negative impact on cell fitness and OLA production ^21^. Thus, *CsAAE1* was expressed in conjunction with *CsOLS* and *CsOAC* in *Y. lipolytica* supplied with 0.5 mM, 1 mM, and 2 mM hexanoic acid at 24 h, respectively. However, we found that there was a decline in the titer of OLA with the overexpression of *CsAAE1* under these conditions when compared to the control strain with no supplementation of hexanoic acid (Figure 3a). The OD_600_ values tested in the process of fermentation showed that the yeast cells grew slowly with supplementation of hexanoic acid at 24 h and their OD_600_ values were lower than that of the control strain at 96 h (Figure 3b). This was probably due to the toxicity of hexanoic acid to *Y. lipolytica* cells. We speculated that only when there were enough yeast cells with *CsAAE1* overexpression, hexanoic acid could be efficiently converted to hexanoyl-CoA, and thus its toxicity to cells could be minimized. Thus, different dosages of hexanoic acid were fed at 48 h. It was found that the titer of OLA was highest with supplementation of 0.5 mM hexanoic acid at 48 h and slightly increased to 0.13 mg/L (Figure 3a), and the yeast cells didn’t have observable changes in the OD_600_ values of the control strain (Figure 3c). Thus, the supplementation of 0.5 mM hexanoic acid at 48 hours was used for further strain screening.

**Figure 3.**
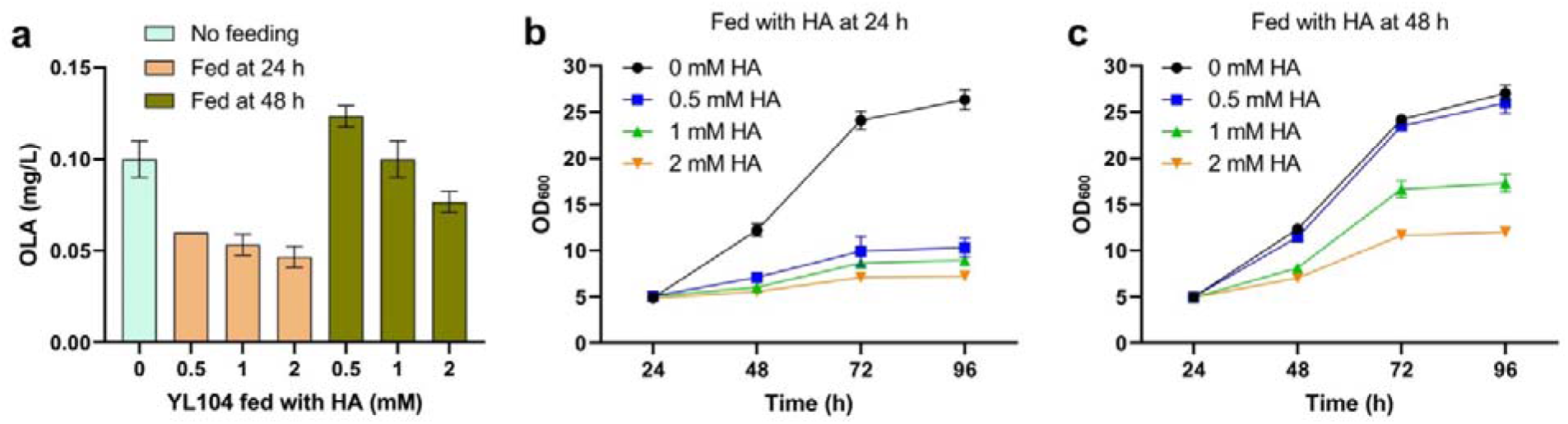
Olivetolic acid (OLA) and cell growth profile of YL104 strain expressing CsAAE1 with the supplementation of hexanoic acid (HA). (**a**) OLA titers in YL104 strain supplied with 0.5 mM, 1 mM, and 2 mM hexanoic acid at 24 h or 48 h. (**b**) Cell growth profile of YL104 strain fed with different dosages of HA at 24 h. (**c**) Cell growth profile of YL104 strain fed with different dosages of HA at 48 h.

Nevertheless, *Cs*AAE1 showed lower catalytic capacity in *Y. lipolytica* than in *S. cerevisiae* ^22^. In order to optimize the conversion of hexanoic acid to hexanoyl-CoA, we screened a panel of six enzymes that have been annotated with acyl-CoA synthase activity. *CsAAE3* encoding a peroxisomal enzyme can accept a variety of fatty acid as substrates including hexanoic acid. We next removed the C-terminal peroxisome targeting sequence 1 (PTS1) and overexpressed the codon-optimized *CsAAE3* in *Y. lipolytica* ^23-25^. Both the short-chain and long-chain fatty acyl-CoA synthetases, *Ec*FadK and *Ec*FadD from *E. coli*, have been shown to exhibit a catalytic activity on C6-C8 fatty acids ^21^. The medium-chain fatty acyl-CoA synthetase *Sc*FAA2 from *S. cerevisiae* and the *Y. lipolytica* native fatty acyl-CoA synthetase encoded by *ylFAA1* (YALI0D17864g) were also tested in this study. The *lvaE* gene of *Pseudomonas putida* KT2440 (*PpLvaE*) encodes an enzyme that is active with C4-C6 carboxylic acids, including hexanoic acid ^26^. Among these chosen hexanoyl-CoA synthetases, we found the overexpression of *PpIvaE* in combination with *CsOLS* and *CsOAC* (strain YL110) resulted in the highest OLA production, which showed an eightfold increase in OLA titer (1.07 mg/L) (Figure 4a). *P. putida* has been reported to exhibit better solvent tolerance and grow under a number of harsh or hydrophobic conditions. We believe that the source of hexanoic acyl-CoA synthase played a major role to unlock the rate-limiting reactions of the initial steps of the cannabinoids pathway. Future enzymology study or directed evolution of *Pp*IvaE may further improve its catalytic function.

**Figure 4.**
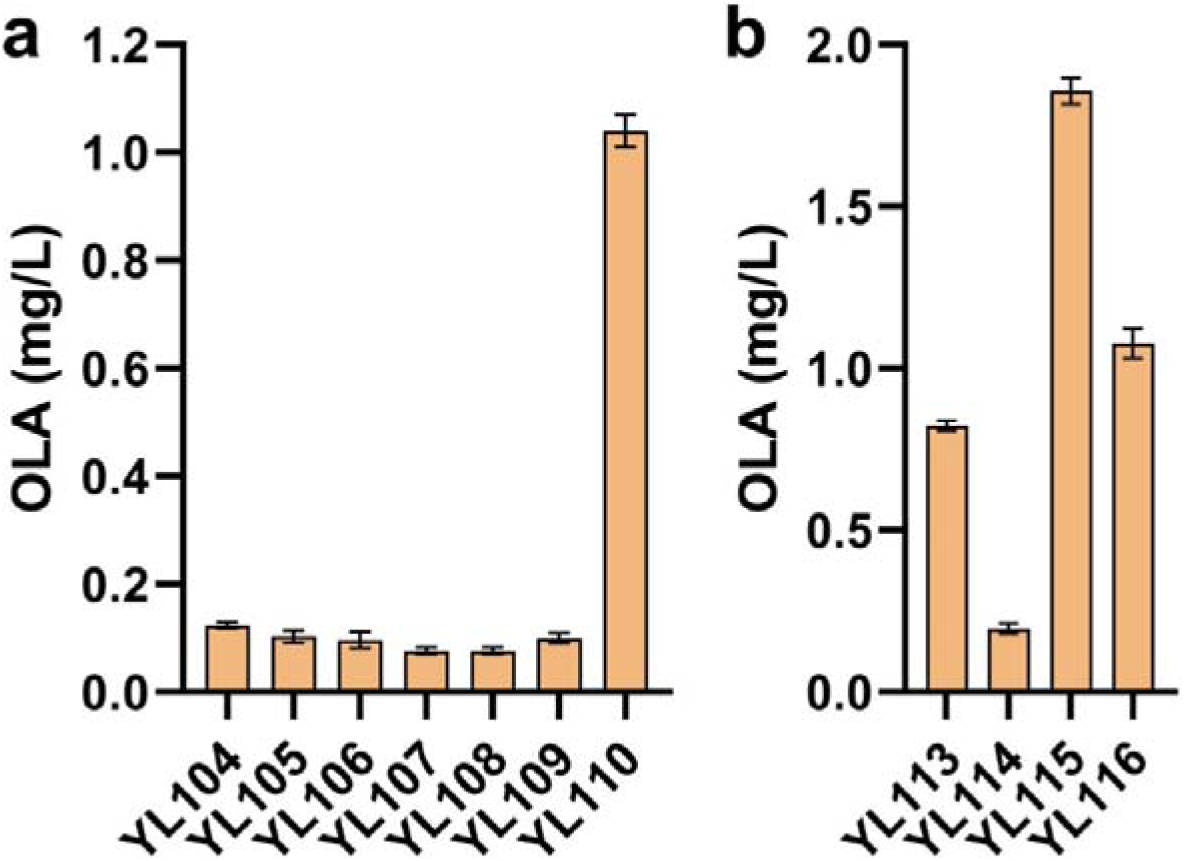
Boosting the supply of precursors hexanoyl-CoA and malonyl-CoA to improve olivetolic acid (OLA) production. (**a**) OLA titers in engineered strains by expressing different hexanoyl-CoA synthetases for optimizing the conversion of hexanoic acid to hexanoyl-CoA. (**b**) OLA titers in engineered strains for enhancing the availability of malonyl-CoA.

### 3.3 Enhancing the availability of precursor malonyl-CoA and controlling pH to improve olivetolic acid production

Malonyl-CoA has been reported as the major rate-limiting precursors for polyketides synthesis ^27^. Next, we used two strategies to improve malonyl-CoA supply. In the first strategy, two endogenous genes *ylDGA1* (YALI0E32769g) and *ylDGA2* (YALI0D07986g) encoding diacylglycerol acyltransferases were knocked out, in such way, we could minimize the formation of triglycerides by blocking the acyltransferase activity of DGA1 and DGA2 ^28, 29^. Increased acyl-CoAs will reduce fatty acids synthase activity and boost the intracellular malonyl-CoA. The plasmid pYLXP’-*CsOLS*-*CsOAC*-*PpLvaE* was transformed into the strain YL111 (*ylDGA2* knockout strain) and YL112 (*ylDGA1* and *ylDGA2* double knockout strain), respectively. Surprisingly, the resulting strains YL113 and YL114 showed a dramatic decline in OLA production, which was contractionary to our expectation (Figure 4b). We speculate that that the increased acyl-CoAs, or acyl-ACPs may strongly feedback inhibit the activity of the acetyl-CoA carboxylase ^30, 31^, which has been reported in both prokaryotic and eukaryotic cells.

In the second strategy, the endogenous gene *ylACC1* (YALI0C11407g) encoding acetyl-CoA carboxylase was overexpressed to facilitate the conversion of acetyl-CoA to malonyl-CoA ^32^. To minimize spliceosomal or transcriptional level regulation, the two internal introns of the native gene *ylACC1* were removed. The engineered strain YL115 with overexpression of ylACC1 produced 1.9 mg/L of OLA, with a 1.8-fold improvement (Figure 4b). Additionally, snf1 has been reported as the protein level regulator for ACC activity. The phosphorylation of the serine groups on ACC will strongly inhibit ACC activity ^33, 34^. For example, a snf1-resistant version of ACC mutant (*ScACC1*^*S659A, S1157A*^) has been reported to boost intracellular malonyl-CoA level and improve 3-hydroxyl propionate or fatty acids production in *S. cerevisiae* ^35^. Thus, a mutant version of *ylACC1*^*S667A, S1178A*^ was obtained after alignment with amino acid sequences of *ScACC1*. However, the result showed that the OLA production failed to achieve improvement by overexpressing *ylACC1*^*S667A, S1178A*^ (strain YL116, Figure 4b). We predict that the phosphorylation modification of ACC1 in *Y. lipolytica* may subject to some hidden layer of regulations, yet to be explored.

We also observed that the pH of the fermentation media dramatically dropped to below 3.5 during the fermentation process, due to the accumulation of organic acids. This decreased pH or increased proton concentration may negatively affect membrane permeability and strain physiology ^32, 36^. We next sought to control the medium pH by using either PBS buffer or CaCO_3_ ^36^. Supplementation of 20 g/L CaCO_3_ maintained stable pH and increased OLA titer by threefold, reaching 5.86 mg/L at 96 h, whereas PBS failed to improve OLA production (Figure 5a). We speculate that the use of PBS buffer may shift the cell metabolism, for example, the cell may increase the phospholipid or cell membrane synthesis ^37^, which is competing with our goal to synthesize OLA because both pathways share the same precursor malonyl-CoA.

**Figure 5.**
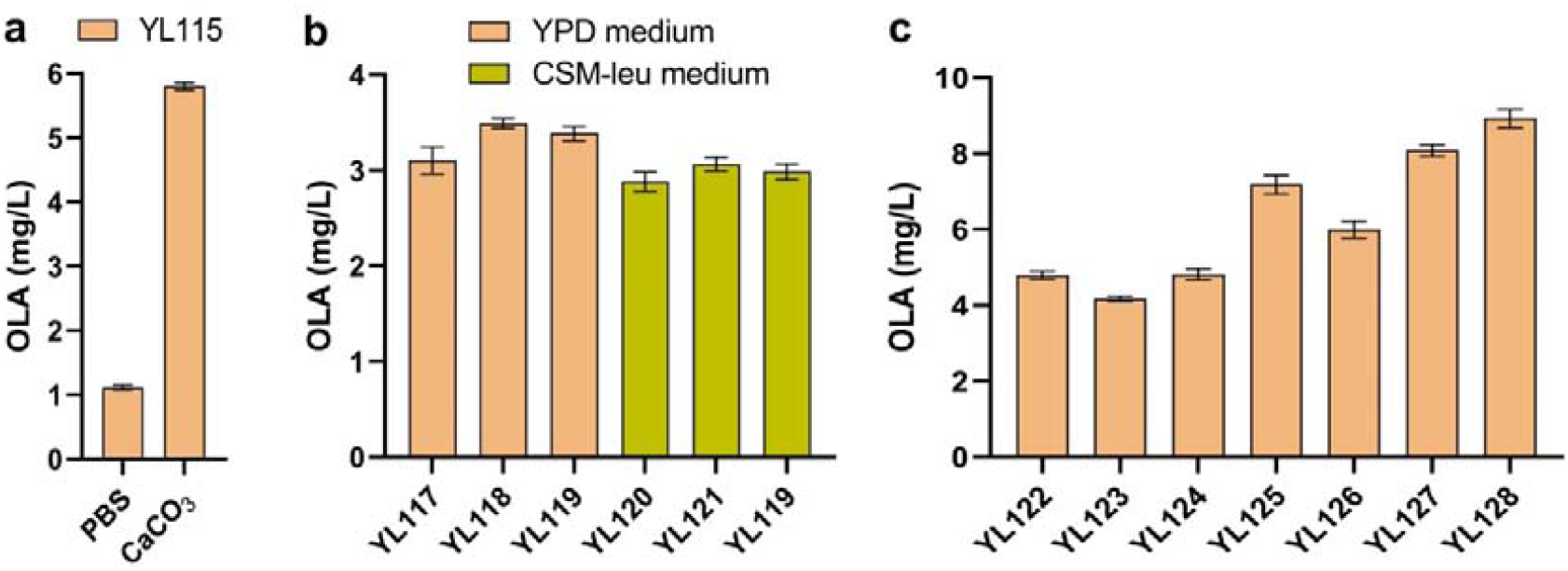
Improve olivetolic acid (OLA) production by controlling pH and increasing acetyl-CoA, ATP and NADPH supply. (**a**) OLA titers after controlling pH of the fermentation medium by using either PBS buffer or CaCO_3_. (**b**) OLA titers in strains of integrating plasmid into different the genomic integration sites in YPD or CSM-leu medium. (**c**) OLA titers in engineered strains by expressing different enzymes and their combinations for improving the supply of acetyl-CoA, ATP and NADPH.

### 3.4 Debottlenecking acetyl-CoA, ATP and NADPH supply to improve olivetolic acid production

Acetyl-CoA served as the basic building block for both hexanoyl-CoA and malonyl-CoA, and the biosynthesis of OLA from acetyl-CoA requires extensive consumption of ATP and NADPH as cofactors to facilitate the activity of ACC, hexanoyl-CoA synthase and olivetolic acid synthase ^21,, 22, 38^ To simplify genetic manipulations and increase strain stability, we next integrated the relevant genes carried by the plasmid pURLB-*CsOLS*-*CsOAC*-*PpLvaE*-*ylACC1* at the genomic locus of YALI0C05907g, which was screened as an orthogonal integration site for polyunsaturated fatty acid production in *Y. lipolytica* ^39^. The integrated strain YL117 yielded about 3.26 mg/L OLA in YPD complex media, which was lower than the production level (5.86 mg/L) of the strain YL115 with chemically-defined complete synthetic media (CSM-leu), possibly due to the altered gene expression profile resulting from genomic integration or shifting of culture media from CSM to YPD (Figure 5b). To probe the effect, YL117 strain was transformed with the empty plasmid pYLXP’ and fermented in CSM-leu media, but the OLA titer decreased to 2.99 mg/L, which indicates YPD media was better for the integrated strain (Figure 5b). Subsequently, the genomic locus of *ku70* and the pBR docking site ^16^ were chosen as integration site for comparison with the genomic locus of YALI0C05907g, resulting in the integrated strains YL118 and YL119, respectively. The strain YL118 in YPD media produced a comparatively higher OLA titer of 3.54 mg/L (Figure 11). By comparisons of the OLA titer from the integrated strain, we speculated that the supply of acetyl-CoA may create a major bottleneck for OLA synthesis in *Y. lipolytica*.

To improve acetyl-CoA supply, the *Y. lipolytica* peroxisomal matrix protein Pex10 (*yl*Pex10) ^40^, *Salmonella enterica* acetyl-CoA synthetase mutant (*Se*Acs^L641P^) ^41^, and *E. coli* pyruvate dehydrogenase complex (*Ec*PDH) with the lipoate-protein ligase A (*Ec*LplA) ^32^ were reported to contribute to boost the availability of acetyl-CoA in *Y. lipolytica* or *S. cerevisiae*. Thus, we overexpressed these enzymes and their combination in the strain YL118. The resultant strains improved the OLA production to different extents, with the strain YL125 overexpressing the combination of ylPex10, *Se*Acs^L641P^ and *Ec*PDH yielding the highest OLA titer of 7.43 mg/L (Figure 5c). Ant1p, a peroxisomal adenine nucleotide transporter, is an integral protein of the peroxisomal membrane and is responsible for transferring ATP into peroxisomes for β-oxidation of medium-chain fatty acids, which will increase the rate of β-oxidation cycle and provide cytosolic acetyl-CoAs ^42^. For example, the *ScANT1* has been overexpressed in *S. cerevisiae* to supply ATP and enhance peroxisome functions ^43^. Thus, we chose to overexpress the *ScANT1* homologous gene *ylANT1* (YALI0E03058g) encoding *Y. lipolytica* peroxisomal ATP/AMP transporter to increase ATP supply and the peroxisomal β-oxidation rate. The native gene *ylMAE1* (YALI0E18634g) encoding malic enzyme (MAE) and *McMAE2* from *Mucor circinelloides* were reported to provide NADPH in *Y. lipolytica* ^32,44^. We next transformed the plasmid pYLXP’-*ylMAE1*-*ylANT1*-*McMAE2* into the strain YL118 to improve the supply for both ATP and NADPH. The resultant strain YL126 produced 6.21 mg/L of OLA (Figure 5c). Next, the expression plasmid containing seven genes which overcome precursor limitations for acetyl-CoA, NADPHs and ATPs were constructed and then transformed into YL118 for coexpression. The resulting strain YL127 further enhanced OLA production to 8.23 mg/L (Figure 5c). Finally, the linearized gene fragments containing pYLXP’-*ylPEX10*-*SeACS*^*L641P*^-*EcPDH*-*EcLplA*-*ylMAE1*-*ylANT1*-*McMAE2* was integrated at the pBR docking site of the strain YL118. The engineered strain YL128 cultured in YPD media produced OLA with a titer of 9.18 mg/L (Figure 5c), which was improved by 83 times as compared to the OLA titer produced by the initial strain YL101 (0.11 mg/L). The current OLA production titer represents a threefold higher than the reported OLA titer produced in engineered *S. cerevisiae* in the shaker flasks ^22^.

## 4. Conclusions

By estimation, more than 75% of the essential drugs are derived from plant secondary metabolites. In their native hosts, the active ingredients of natural products are in extremely low concentration (most often < 0.01%) and subject to environmental, seasonal, and regional variations ^45^. Current plant breeding and agricultural technologies cannot meet the market demand. Total chemical synthesis is not economical feasible because of the structural complexity and existence of chemical analogs. Engineering plant biosynthetic pathways in promising microbial hosts offers significant promise for scalable synthesis of essential drug components.

In this study, we have systematically investigated the enzymatic bottlenecks that restrain the efficient biosynthesis of CBD precursor olivetolic acid in the oleaginous yeast *Y. lipolytica*. We took a reverse engineering approach and sequentially identified that the supply of hexanoyl-CoA, malonyl-CoA, acetyl-CoA, NADPH and ATP are the rate-limiting steps. To overcome these limitations, we have screened enzymes aiming to debottleneck the pathway limitations and redirect the carbon flux toward the end-product olivetolic acid. We discovered that the use of the *P. putia* IvaE encoding a novel acyl-CoA synthase could efficiently convert exogenously-added hexanoic acid to hexanoyl-CoA. The co-expression of the acetyl-CoA carboxylase, the pyruvate dehydrogenase bypass, the NADPH-generating malic enzyme, as well as the activation of peroxisomal β-oxidation pathway and ATP export pathway were efficient strategies to remove the pathway bottlenecks. Collectively, these strategies have led us to construct a Y. *lipolytica* strain that produced olivetolic acid at a titer 83-fold higher than our initial strain (0.11 mg/L). While the current production level is low, we expect these strategies may serve as a baseline for other metabolic engineers who are interested in engineering CBD biosynthesis in oleaginous yeast species.

## Acknowledgement

This project is supported by the Bill & Melinda Gates Foundation Award (*grant number OPP 1188443) and National Science Foundation (CBET-1805139)*. We thank Dr. Joshua Wilhide and Dr. LaCourse to help run the LC-MS characterization of OLA at the UMBC Molecular Characterization and Analysis Complex center (UMBC chemistry facility) and help prepare the LC-MS report.

## Author contributions

PX designed the study, analyzed the data and revised the manuscript. J.M. performed the genetic engineering, enzyme screening and cell cultivation with input from YG. J.M wrote the manuscript and analyzed the data.

