## Supplementary tables and figures for "Biosynthesis of cannabinoid precursor olivetolic acid by overcoming rate-limiting steps in genetically engineered *Yarrowia lipolytica*"

**Table 1. Primers used in this study**

| Primers | Sequence |
|---|---|
| CsOAC-F | ccgaccagcactttttgcagtactaaccgcaggccgtgaaacaccttattgtcctc |
| CsOAC-R | ggacaggccatggaactagtcggtaccttatttacgaggggtgtagtcgaaaataag |
| CsOLS-F | ccgaccagcactttttgcagtactaaccgcagaatcacctccgggcagaagg |
| CsOLS-R | ggacaggccatggaactagtcggtaccttaatacttaatggggacgcttc |
| CsOAC-Gly-F | ggaggaggtggtggtggtggtggaggtggcatggccgtgaaacaccttattgtcctc |
| CsOLS-Gly-R | acggccatgccacctccaccaccaccaccacctcctccatacttaatggggacgcttct |
| EcFadD-F | ccgaccagcactttttgcagtactaaccgcagaagaaggtttggcttaaccgttatcc |
| EcFadD-R | ggacaggccatggaactagtcggtacctcaggctttattgtccactttgccgcgcg |
| EcFadK-F | tccgaccagcactttttgcagtactaaccgcagcatcccacaggcccgcatctcgg |
| EcFadK-R | ggacaggccatggaactagtcggtaccttattcaatctcttcacagacatc |
| ScFAA2-F | tccgaccagcactttttgcagtactaaccgcaggccgctccagattatgcacttaccg |
| ScFAA2-R | ggacaggccatggaactagtcggtaccctaaagcttttctgtcttgactag |
| ylACC1-F | tccgaccagcactttttgcagtactaaccgcagcgactgcaattgaggacactaacacg |
| ylACC1-R | ggacaggccatggaactagtcggtacctcacaacccccttgagcagctcagcccg |
| ylACC1(S667A)-F | agttagacctcttgctgacggtggtattc |
| ylACC1(S667A)-R | aataccaccgtcagcaagaggtctaactc |
| ylACC1(S1178A)-F | gtctcgagctgatgccgtctccgacttttc |
| ylACC1(S1178A)-R | aagtcggagacggcatcagctcgagacac |
| ylFAA1-F | ccgaccagcactttttgcagtactaaccgcaggtcggatacacaatttcctcaaagcc |
| ylFAA1-R | ggacaggccatggaactagtcggtaccctaagactgctcgtagcactcatcaatttc |
| pTEF-Rvs | ccggatggccagacaaagaaaca |
| XPR2-Fw | gtaaatagaaaatctggcttgtaggtggcaaaat |
| Ku70_DwF | gtccggagcggccgcGCATGCaagtcgacaCTAGGGAGGCACATCTAAACGAATAACG |
| Ku70_DwR | gttacatccttttatcagacatacctaggAGTGAACGACCAAGACTAAAGGGTG |
| ku70_UpF | ccctaaatttgatgaaagcctaggCGACTTGATGTTTAGAGTGTCCAGATCC |
| ku70_UpR | taatgtatgctatacgaagttatTTTCAAAAAGCGGCGGTTCGTG |
| ku70_ChkF | GCCAAGTTCTCTTTCCCCTACATG |
| ku70_ChkR | CTTCAGTAACTGGGCCCACGC |
| YliC05907-UpF | tggcatccctaaatttgatgaaagcctaggACAGTTCTCTTCTCCTTGTTGAGATATAC |
| YliC05907-UpR | acttcgtataatgtatgctatacgaagttatGGTGGAGTCATGTGAATTGAGCGCGGTG |
| YliC05907-DwF | cgtccggagcggccgcGCATGCaagtcgacAGAGTATAGTAATGTATTATTTGCTTAGG |
| YliC05907-DwR | cctatgttacatccttttatcagacatacctaggCCGAACCAAGGAGATGATCAAGTCC |
| YliC05907-ChkF | ATTGCCGAGATTTCCGCAAAAACCTGAAG |
| YliC05907-ChkR | AAGAACAAGACCGACAACCTGCCTGTTGG |
| YliDGA1_DwF | gctagcgagacaataacggaggaGGAAAACTGCCTGGGTTAGGCAAAT |
| YliDGA1_DwR | atcagacatagcggccgcTCTCTGATGGCCTGGAGCGAG |
| YliDGA1_UpF | aatttgatgaaaggcggccgcATGCTGCGGGCGGATCCTGG |
| YliDGA1_UpR | tgtatgctatacgaagttatAGCTTTTGTTTTGTGTGACTTGTCTGT |
| YliDGA1_ChkF | GTTTATGCATTCTGTTGGACCTTAGTCTG |
| YliDGA1_ChkR | GTTATCTACCACGATTTTTTGGTTTCTGAGGC |

| | |
|---|---|
| YliDGA2_DwF | gctagcgagacaataacggaggaCATAACACTCATCAGTAGCCTTTACAGTGAT |
| YliDGA2_DwR | ccttttatcagacatagcggccgcTTGCTCTTGTAATTCCATAGATAATATATACGAAA |
| YliDGA2_UpF | aaatttgatgaaaggcggccgcCTTGGGAGTGTATTTGGAAAATGACTTGG |
| YliDGA2_UpR | tgtatgctatacgaagttatTTTGCGGGCGGTACGGGTACA |
| YliDGA2_ChkF | CTATCGCCCCAAAGTGTTTCTAGCA |
| YliDGA2_ChkR | GAGATGGCATGCCAACGTTGAC |
| ylPEX10-F | tccgaccagcactttttgcagtactaaccgcagtggggaagttcacatgcattcgctgg |
| ylPEX10-R | ggacaggccatggaactagtcggtaccttatctgataggcaacaagttctgctctc |
| EcLpdA-F | ccgaccagcactttttgcagtactaaccgcagagtactgaaatcaaaactcaggtc |
| EcLpdA-R | GGACAGGCCATGGAACTAGTCGGTACCTTACTTCTTCTTCGCTTTCGGGT |
| EcAceF-F | ccgaccagcactttttgcagtactaaccgcaggctatcgaaatcaaagtaccggac |
| EcAceF-R | GGACAGGCCATGGAACTAGTCGGTACCTTACATCACCAGACGGCGAATGTC |
| EcAceE-F | ccgaccagcactttttgcagtactaaccgcagtcagaacgtttcccaaatgacgtg |
| EcAceE-R | ggacaggccatggaactagtcggtaccttacgccagacgcgggttaactt |
| EcLplA-F | ccgaccagcactttttgcagtactaaccgcagtccacattacgcctgctcatctc |
| EcLplA-R | GGACAGGCCATGGAACTAGTCGGTACCCTACCTTACAGCCCCCGCCATCCATG |
| ylMAE1-F | ccgaccagcactttttgcagtactaaccgcagttacgactacgaaccatgcgaccc |
| ylMAE1-R | GGACAGGCCATGGAACTAGTCGGTACCCTAGTCGTAATCCCGCACATGGATG |
| ylANT1-F | tccgaccagcactttttgcagtactaaccgcaggcagctatttccaaagactatgttc |
| ylANT1-R | ggacaggccatggaactagtcggtaccttatcccttgatcaaggtgggggccttcatg |

**Table 2. Plasmids used in this study**

| Plasmids | Characteristics |
|---|---|
| pYLXP' | pYaliA1 vector backbone with leucine marker and Ampicillin resistance gene |
| pYLXP'-CsOLS | pYLXP' containing gene CsOLS |
| pYLXP'-CsOAC | pYLXP' containing gene CsOAC |
| pYLXP'-CsOLS-CsOAC | pYLXP' containing gene CsOLS and CsOAC |
| pYLXP'-CsOLS-Gly-CsOAC | pYLXP' containing gene CsOLS and CsOAC with Glycine linker |
| pYLXP'-CsOAC-Gly-CsOLS | pYLXP' containing gene CsOAC and CsOLS with Glycine linker |
| pYLXP'-CsAAE1 | pYLXP' containing gene CsAAE1 |
| pYLXP'-CsAAE3 | pYLXP' containing gene CsAAE3 |
| pYLXP'-EcFadD | pYLXP' containing gene EcFadD |
| pYLXP'-EcFadK | pYLXP' containing gene EcFadK |
| pYLXP'-ScFAA2 | pYLXP' containing gene ScFAA2 |
| pYLXP'-ylFAA1 | pYLXP' containing gene ylFAA1 |
| pYLXP'-PpLvaE | pYLXP' containing gene PpLvaE |
| pYLXP'-CsOLS-CsOAC-CsAAE1 | pYLXP' containing gene CsOLS, CsOAC and CsAAE1 |
| pYLXP'-CsOLS-CsOAC-CsAAE3 | pYLXP' containing gene CsOLS, CsOAC and CsAAE3 |
| pYLXP'-CsOLS-CsOAC-EcFadD | pYLXP' containing gene CsOLS, CsOAC and EcFadD |
| pYLXP'-CsOLS-CsOAC-EcFadK | pYLXP' containing gene CsOLS, CsOAC and EcFadK |
| pYLXP'-CsOLS-CsOAC-ScFAA2 | pYLXP' containing gene CsOLS, CsOAC and ScFAA2 |
| pYLXP'-CsOLS-CsOAC-ylFAA1 | pYLXP' containing gene CsOLS, CsOAC and ylFAA1 |
| pYLXP'-CsOLS-CsOAC-PpLvaE | pYLXP' containing gene CsOLS, CsOAC and PpLvaE |
| pYLXP'-ylACC1 | pYLXP' containing gene ylACC1 |
| pYLXP'-ylACC1$^{S667A}$ | pYLXP' containing gene ylACC1$^{S667A}$ |
| pYLXP'-ylACC1$^{S667A,S1178A}$ | pYLXP' containing gene ylACC1$^{S667A,S1178A}$ |
| pYLXP'-CsOLS-CsOAC-PpLvaE-ylACC1 | pYLXP' containing gene CsOLS, CsOAC, PpLvaE and ylACC1 |
| pYLXP'-CsOLS-CsOAC-PpLvaE-ylACC1$^{S667A,S1178A}$ | pYLXP' containing gene CsOLS, CsOAC, PpLvaE and ylACC1$^{S667A,S1178A}$ |
| pYLXP'-ylPEX10 | pYLXP' containing gene ylPEX10 |
| pYLXP'-SeACS$^{L641P}$ | pYLXP' containing gene SeACS$^{L641P}$ |
| pYLXP'-EcPDH-EcLplA | pYLXP' containing gene EcPDH and EcLplA |
| pYLXP'-ylPEX10-SeACS$^{L641P}$-EcPDH-EcLplA | pYLXP' containing gene ylPEX10, SeACS$^{L641P}$, EcPDH and EcLplA |
| pYLXP'-ylMAE1-ylANT1-McMAE2 | pYLXP' containing gene ylMAE1, ylANT1 and McMAE2 |
| pYLXP'-ylPEX10-SeACS$^{L641P}$-EcPDH-EcLplA-ylMAE1-ylANT1-McMAE2 | pYLXP' containing gene ylPEX10, SeACS$^{L641P}$, EcPDH, EcLplA, ylMAE1, ylANT1 and McMAE2 |
| pYLXP'-loxP-ura | pYLXP'containing the loxP-URA-loxP cassette |
| pYLXP'-loxP-hygr | pYLXP'containing the loxP-hygr-loxP cassette |
| pYLXP'-Cre | pYLXP'containing gene Cre |
| pURLA | *Ku70* site integration plasmid |
| pURLB | *YALI0C05907g* site integration plasmid |

| | |
|---|---|
| pYLXP'-loxP-ura-ylDGA2 | pYLXP'-loxP-ura containing gene ylDGA2 deletion cassette |
| pYLXP'-loxP-hygr-ylDGA1 | pYLXP'-loxP-hygr containing gene ylDGA1 deletion cassette |
| pURLA-CsOLS-CsOAC-PpLvaE-ylACC1 | pURLA containing gene CsOLS, CsOAC, PpLvaE and ylACC1 |
| pURLB-CsOLS-CsOAC-PpLvaE-ylACC1 | pURLB containing gene CsOLS, CsOAC, PpLvaE and ylACC1 |

NOTE: Cs, *Cannabis sativa*; Ec, *Escherichia coli*; Sc, *Saccharomyces cerevisiae*; yl, *Yarrowia lipolytica*; Pp, *Pseudomonas putida* KT2440; Se, *Salmonella enterica*; Mc, *Mucor circinelloides*.

**Table 3. Strains used in this study**

| Strains | Characteristics |
|---|---|
| po1g | Wild-type strain W29 (ATCC20460) derivate; W29 ΔmatAΔxpr2-332 Δaxp-2 Δleu2-270 pBR platform |
| po1f | po1g derivate; Further deletion of gene ura; po1g Δura3 |
| po1fk | po1f derivate; Further deletion of gene ku70; po1f Δku70::loxP |
| YL100 | po1fk with the empty plasmid pYLXP’ |
| YL101 | po1fk with plasmid pYLXP'-CsOLS-CsOAC |
| YL102 | po1fk with plasmid pYLXP'-CsOLS-Gly-CsOAC |
| YL103 | po1fk with plasmid pYLXP'-CsOAC-Gly-CsOLS |
| YL104 | po1fk with plasmid pYLXP'-CsOLS-CsOAC-CsAAE1 |
| YL105 | po1fk with plasmid pYLXP'-CsOLS-CsOAC-CsAAE3 |
| YL106 | po1fk with plasmid pYLXP'-CsOLS-CsOAC-EcFadD |
| YL107 | po1fk with plasmid pYLXP'-CsOLS-CsOAC-EcFadK |
| YL108 | po1fk with plasmid pYLXP'-CsOLS-CsOAC-ScFAA2 |
| YL109 | po1fk with plasmid pYLXP'-CsOLS-CsOAC-ylFAA1 |
| YL110 | po1fk with plasmid pYLXP'-CsOLS-CsOAC-PpLvaE |
| YL111 | po1fk derivate; Further deletion of genes ylDGA2; po1fk ΔylDGA2::loxP |
| YL112 | YL111 derivate; Further deletion of genes ylDGA1; po1fk ΔylDGA2ΔylDGA1::loxP |
| YL113 | YL111 with plasmid pYLXP'-CsOLS-CsOAC-PpLvaE |
| YL114 | YL112 with plasmid pYLXP'-CsOLS-CsOAC-PpLvaE |
| YL115 | po1fk with plasmid pYLXP'-CsOLS-CsOAC-PpLvaE-ylACC1 |
| YL116 | po1fk with plasmid pYLXP'-CsOLS-CsOAC-PpLvaE-ylACC1$^{S667A,S1178A}$ |
| YL117 | po1fk derivate; Further integration of genes CsOLS-CsOAC-PpLvaE-ylACC1 at *YALI0C05907g* site; po1fk CsOLSCsOACPpLvaEylACC1::loxP |
| YL118 | po1fk derivate; Further integration of genes CsOLS-CsOAC-PpLvaE-ylACC1 at *ku70* site; po1fk CsOLSCsOACPpLvaEylACC1::loxP |
| YL119 | po1fk derivate; Further integration of genes CsOLS-CsOAC-PpLvaE-ylACC1 at pBR platform; po1fk CsOLSCsOACPpLvaEylACC1::Leu |
| YL120 | YL117 with the empty plasmid pYLXP’ |
| YL121 | YL118 with the empty plasmid pYLXP’ |
| YL122 | YL118 with plasmid pYLXP'-ylPEX10 |
| YL123 | YL118 with plasmid pYLXP'-SeACS$^{L641P}$ |
| YL124 | YL118 with plasmid pYLXP'-EcPDH-EcLplA |
| YL125 | YL118 with plasmid pYLXP'-ylPEX10-SeACS$^{L641P}$-EcPDH-EcLplA |
| YL126 | YL118 with plasmid pYLXP'-ylMAE1-ylANT1-McMAE2 |
| YL127 | YL118 with plasmid pYLXP'-ylPEX10-SeACS$^{L641P}$-EcPDH-EcLplA-ylMAE1-ylANT1-McMAE2 |
| YL128 | YL118 derivate; Further integration of genes ylPEX10-SeACS$^{L641P}$-EcPDH-EcLplA-ylMAE1-ylANT1-McMAE2 at pBR platform; YL118 ylPEX10SeACS$^{L641P}$EcPDHEcLplAylMAE1ylANT1McMAE2::Leu |

**Codon-Optimized gene sequences**

>CsOLS

atgaatcacctccgggcagaaggccccgcatccgtcctggctatcggtactgcaaatcccgaaaacatccttctccaagacgagttccccgattat
tatttccgagtcaccaagtctgagcacatgactcaactcaaggagaaattccggaagatttgtgacaaaagcatgatccgtaagcggaactgtttcc
tgaatgaagagcacctgaagcagaatcctcgacttgttgaacacgagatgcagacccttgatgcacgtcaggacatgcttgttgtggaggttccca
agctgggtaaagatgcctgtgccaaggctatcaaggagtggggccagcctaaatcgaagatcacgcatctgattttcacctctgcctcgacaactg
atatgcccggagctgattatcactgtgcaaaactgctcggtctgagcccctctgtcaaacgggttatgatgtaccagctgggttgctacggcggtgg
aacggtgctgcgtatcgcaaaggatattgccgaaaataacaaaggcgccagagtccttgctgtgtgttgtgacattatggcctgtctcttccgggga
ccctccgaatccgatcttgaactgcttgtcggacaagcaatttttggtgatggcgcagccgccgttatcgttggtgcagagcctgatgaatctgtgg
gtgaaagacccattttcgagctcgtctcgacaggtcaaactatcctgcccaattcggagggcacaattggtggacatattcgtgaagcaggactga
ttttcgacctgcacaaagatgttcctatgcttatctccaataacatcgagaaatgccttatcgaggcctttaccccctatcggcatttcggattggaattcg
attttttggatcacccatcccggaggaaaagcaatcctcgataaagtggaagaaaaacttcacctcaagtccgataagttcgtggactcccggcatg
tgctctctgaacatggaaatatgtcgtcctctactgtccttttcgtgatggacgaacttcggaagcggtccctcgaggaaggcaaaagcacaacag
gtgacggatttgaatggggtgttctctttggctttggccctggccttacggttgaacgagtcgttgttagaagcgtccccattaagtattaa

>CsOAC

atggccgtgaaacaccttattgtcctcaaattcaaggacgaaatcactgaggcacagaaggaggaatttttaaaacctacgtcaatctcgtcaatat
cattcctgctatgaaggatgtctattggggcaaagatgtgacccaaaaaaacaaggaagaaggatacacccacattgtcgaggttacattcgaga
gcgttgaaacgattcaagactacatcattcaccccgctcacgttggctttggtgacgtttaccgatccttttgggaaaagctccttattttcgactacac
ccctcgtaaa

>CsAAE1

atgggaaagaactacaagtcgcttgattccgtggtcgcctcggattttatcgctctgggtattacatcggaagtcgctgagactcttcatggccggct
cgcagaaattgtttgtaactatggagctgccacgcctcagacttggatcaacattgctaatcacattctctcccccgaccttcctttctccctgcatcaa
atgctcttctatggctgctacaaggatttcggtcctgcacctcctgcttggattcctgatcctgagaaagttaaatctactaacctcggtgcccttctgg
agaaacgtggtaaagaattcctgggtgtgaaatacaaggaccctatttcgtccttttctcacttccaagagttctccgttcgaaatcctgaggtctattg
gcgtacggtccttatggatgagatgaagatttctttttccaaagatcctgagtgcattctccgtcgagacgacattaataaccccggtggttccgaatg
gcttcctggtggatatctcaattccgctaaaaattgcctcaatgttaactccaataaaaagctgaatgatacgatgatcgtttggcgagacgaaggca
atgacgacctgccccttaacaaactcactctcgaccagctgcgaaaacgagtctggcttgtgggctacgcccttgaagagatgggtctcgaaaaa
ggatgtgccatcgctattgacatgcccatgcatgttgacgccgtggtgatctacctcgcaattgttcttgctggatacgtggtggtgagcatcgccga
cagcttttcggcacccgagattagcacgcggctcagactctcgaaagccaaagctatcttcacccaagatcatattatccgaggcaaaaaacggat
ccctctctattcgcgtgtggtcgaagcaaaaagccccatggccatcgttattccctgttctggctccaacatcggcgcagagcttagagacggcgat
atctcctgggactacttcctcgagagagctaaggagtttaagaactgtgaattcactgcacgagagcagcccgtcgatgcttacaccaacatcctgt
ttagctctggcaccactggtgaacctaaggcaatcccttggacccaagctacacctcttaaggctgcagcagatggttggtcgcatctcgacatcc
gaaagggcgatgttatcgtgtggcctactaaccttggctggatgatgggtccttggctcgtctatgcctctctgctcaacggtgccagcatcgccctt
tataacggtagccccccttgtctccggcttcgcaaagtttgtgcaagatgccaaggttacaatgcttggtgtcgtcccttctattgttagaagctggaaa
agcacaaattgcgtttccggctacgactggtctactattcggtgctttagcagctcgggtgaagcctcgaatgtggatgaatacctctggctgatgg
gtagagcaaattacaaacctgtcattgagatgtgtggcggaaccgagatcggaggagccttctcggctggatcctttcttcaagctcaatcccttag
ctctttttcttcgcagtgtatgggttgtacgctgtatattcttgataaaaatggataccccatgcctaaaaacaaacctggtattggagagcttgccctg
gccctgtcatgttcggcgcttcgaaaacacttctgaacggcaaccaccacgacgtttatttcaagggtatgcccacgctcaacggtgaagtccttcg
acgtcatggcgatattttcgaactgacaagcaacggatactatcatgctcatggtagagccgatgacacgatgaacattggtggaatcaagatctct
tccatcgagattgaacgggtctgtaacgaagttgacgaccgtgtcttcgaaacgacggccattggagtgcctcctcttggtggtggtcctgagcag

ctcgttattttttcgttctcaaggattctaatgatacaactatcgacctgaaccagctgcggctgtctttcaatcttggactgcaaaaaaagctcaatcc
ccttttcaaagtcacgcgtgtggtgcctctctcgtcgctgcctcggactgccaccaataaaattatgcggcgggtgctccgacagcagttttcccattt
cgagtaa

>CsAAE3
atggagaagagcggttatggacgggatggaatctatcgttctctgcgtcccccccttcatctgcctaataacaataacctgagcatggtgtcgttcct
cttccgaaactcctcttcctatcctcagaaacccgcacttatcgactctgagacgaatcaaattctttccttttcccacttcaagtccacagtcatcaaag
tctcgcacggatttcttaaccttggaattaagaagaacgatgtcgttctgatttatgcccctaattcgatccacttccccgtttgctttctcggtattattgc
atcgggcgctattgccaccacgtccaatcctctctatactgtcagcgaactttctaagcaggtgaaggactcgaatcctaaactcattatcacggtgc
cccaactcctcgaaaaggtgaaaggattcaatctccctaccattctcattggccctgattcggaacaggaatccagcagcgataaggtcatgacatt
taatgacctcgttaaccttggtggctcttccggctctgaatttcccattgtggatgatttcaaacaatctgacacggccgcactcctttactcgagcgga
acaaccggaatgtcgaagggcgtcgttctcacgcacaagaatttcatcgcaagctctcttatggttaccatggagcaagacctggtcggtgaaatg
gacaacgtttttctttgttttcttcccatgttccacgttttcggtctcgctattattacttacgcccaacttcagcgtggaaataccgtcatttctatggccag
atttgatcttgaaaagatgctcaaggatgttgaaaagtacaaggtgactcacctctgggtcgttcctcccgtgatccttgccctttctaagaatagcat
ggttaaaaagtttaacctttctagcattaaatatatcggtagcggcgctgcacctcttggcaaggaccttatggaggaatgcagcaaagtcgtgccct
atggaatcgtggctcaaggctatggcatgaccgagacttgcggtattgtctctatggaagatattcgaggtggaaaacggaattctggtagcgcag
gcatgctcgcttcgggagttgaggctcaaattgtttccgttgataccctgaagcctctccctcctaaccagctcggagaaatttgggtgaaaggacc
caacatgatgcaaggatacttcaacaaccctcaggcaactaagctgacaattgacaagaaaggttgggtgcataccggagatctgggctactttga
cgaggacggacatctctacgtcgttgatcgaatcaaggaactcatcaaatacaagggcttccaagtcgcacccgctgagctcgaaggactgctgg
tcagccaccctgagattctggacgctgtcgtcattcccttccccgatgcagaagccggagaggttcctgttgcctacgtggtcagatcgcccaactc
gtccctgactgaaaatgatgtcaagaaatttatcgctggccaagtcgcttcttttaagcggctgcgaaaggtcactttcattaacagcgttcctaaaag
cgcttctggaaaaattctttaa

>PpLvaE
atgatggtgcccacgcttgagcatgagctggctcctaacgaggctaaccacgtccctctcagccccctttcgttcctgaagcgagcagcacaagt
gtatcctcaacgggatgctgtcatctacggtgcaagacgatactcgtatcgtcagcttcatgaacgttctcgagctcttgcttcggctcttgagcgggt
gggcgtccagcctggagagcgtgtcgctatcctggcccccaacattcctgaaatgctggaagcccattatggcgtgcccggagccggtgcagtg
ctcgtctgtattaacatccgtcttgagggccggtctattgccttcatcctccggcactgtgcagcaaaagtcctcatttgcgaccgtgagtttggtgct
gttgcaaatcaggcactcgctatgctggacgcacctcccctgctggtcggaatcgacgacgaccaagctgaacgggccgatctggcacacgatc
tcgactatgaagccttccttgcacagggagatcctgcacgtcctctgtctgcccccaaaacgagtggcagtctatcgccattaactatacttccgg
aacaacaggtgatcccaaaggagtcgttctgcaccaccgtggagcatatcttaacgcctgtgctggtgccctcatcttccaactgggcccccggtc
tgtgtatctgtggactcttcccatgttccattgcaacggatggtctcatacttgggctgtgactctctcgggcggtacgcacgtttgtcttcgaaaggtg
caacctgacgcaatcaatgccgccattgctgaacacgcagttacccatctgtcggccgcacccgtcgtcatgtccatgctcatccacgccgagca
cgcatcggcacctcccgttcctgtgtccgtgattacgggaggagctgcccccccttcggccgtcattgctgcaatggaagcacggggctttaatatt
actcacgcttacggaatgacagagtcgtatggaccttcgactctctgcctgtggcagcctggtgtcgacgagctgcccctggaagccagagccca
gtttatgtcgagacagggtgtcgctcacccccctcctcgaagaagccaccgtgctcgacacggatacgggacgacccgtccctgcagacggcctg
actctcggtgaactcgttgttcgtggaaacacagtcatgaaaggatacctccataatcccgaagcaactcgtgcagccctcgctaacggttggctg
cacacgggcgatctcgccgtgctccatctcgatggatatgttgaaattaaggatagagcaaaagatattatcatctcgggaggtgagaatatttcctc
gctcgagatcgaagaggttctgtaccagcaccccgaggtggtcgaggctgcagtcgttgccagacctgactctcgttggggcgagacgcccca
cgcattcgtcaccctcagagccgacgcactggcttccggtgacgatctcgtgagatggtgtagagaacgactggcccattttaaagctcctcggca
cgtgtctctggtcgatcttcctaaaacagccaccggaaaaatccaaaaatttgtcctccgtgaatgggcacgacagcaagaggcacagatcgcag
atgctgagcattaa

>SeACS$^{L641P}$
atgagccaaactcacaagcatgcaatccccgccaacattgccgaccgttgcctcattaaccctgaacagtacgaaactaagtacaaacaatccatc
aacgatcctgataccttttggggagagcagggtaaaattctggattggattacaccttatcagaaagtgaaaaatacatcttttgcccctggcaatgtc
tccatcaaatggtatgaagacggcaccctgaatctggctgctaattgtcttgaccgacacctgcaagaaaacggagatcggactgctattatttggg
aaggagatgacacgagccagtccaagcatatctcgtatcgggagctgcatcgagacgtgtgtcggtttgccaatactctcctcgaccttggaatta
aaaaaggtgatgtggtggccatctatatgcctatggttcctgaggccgctgtggctatgctggcctgtgcccgtattggtgctgtgcatagcgtcattt
ttggaggcttctcgcctgaggcagtcgcaggacggatcattgattcctcctcccggctggtgatcactgcagacgaaggagtccgggctggacg
atctattcccctgaagaagaatgtggacgacgcactcaagaaccctaacgtgacctctgttgagcatgtgatcgtgctgaaacgaacaggctctga
tattgattggcaagagggacgtgatctgtggtggcgtgacctgattgaaaaagcatcgcccgagcaccaacctgaggcaatgaatgcagaagac
cccctgttcatcctgtacacctcgggatctacaggtaaacccaaaggcgtgctccataccacgggtggataccttgtctacgccgcaactacgttca
agtacgtgtttgactatcatcccggagacatttactggtgtacagctgatgttggatgggttacaggtcattcctatcttctgtacggccctctcgcttgt
ggcgcaactacacttatgtttgagggtgttcctaattggcccacacctgctcgaatgtgccaggtggtggataaacaccaagtgaacattctgtaca
cagcccccacagccatccgggccctcatggccgaaggcgacaaagctatcgagggtacggaccgatcttcgctccggatccttggctctgttgg
agagcctatcaatcctgaagcttgggagtggtattggaagaagattggcaaggagaagtgccccgtggttgacacatggtggcagacggagac
gggaggattcatgatcacccctctccccggagccatcgaactgaaagccggaagcgcaactcgtcccttttttggcgttcaacccgcccttgtgga
caacgagggacacccccaggagggcgctacagaaggtaatcttgttatcacagactcctggcccggacaggctcgaacactctttggcgatcac
gaacggtttgagcaaacctacttcagcacatttaaaaacatgtacttttccggagatggtgcacggcgggacgaagacggctactactggattacg
ggaagagttgatgatgttctcaacgtctctggccaccgacttggaacggcagaaatcgaatctgccctcgtcgcccatcctaagattgccgaagcc
gcagttgtcggaattccccacgccatcaaaggtcaagccatctacgcttacgtgactctcaaccatggtgaagaacctagccccgagctgtacgc
cgaggtcagaaactgggttcgaaaggaaattggtcctcttgcaacgcctgatgtccttcattggactgattccctgcctaaaacccggagcggaaa
gattatgcgacgaatccttcggaagatcgccgctggtgacacgtcgaacctcggcgatacctctacactggccgaccccggagtggtcgaaaaa
cctcttgaggagaaacaagcaatcgcaatgccctcctaa

>McMAE2
atgtcgcccatcatcgatttcgtgcgacgtcagcttagctctacaaaacttcacgaggaacagcaaaccgccactacgaacgatctcgtgtcgcgtt
ccggatatcttaacgagggcaaatatgaagttcggcttaactgcatcaacgctggttgtcttcaaaaaaaactcaactatatcggtactgcaatggac
cctgctaaacgtcaaagacttggcctgaatggtcttctccccgccggtgtcgaaacccttgaaattcagaaggcccgagccctgcgtgtcctgcgtt
ccaaacacaatctcctcgagaaatatattcttatggcccaactgcgtacaacgaacgttcggcttttctataagattgttattgacgaactcgaaacgg
tgcaacttgcccccgtcatttacactcccaccgtcggaaccgcttgtctggagtatagcactatctacccctttctcgcagcacctggtgttcccgatg
gactttacctcacaaaggcagagctccccgaactgtgccagacaatccgtaactacagacccactgacactgaaggttttgagcccgagattgcc
gttatcagcgacggttctcggatcctgggtcttggcgacctgggcaccaacggtatgggtattcctatgggtaaacttcaactgtacgtggccggc
gctggaatcgaccctcgacgtacactccccatcattctcgaccttggtaccaataatgaaaaactgctcaacgatgagttttatatcggccttcggca
gaaaagacctaatgacgaagagttttaccagacagttgataccgtgcttactgccctccatactgtgtatcccaacctcctcattcagttcgaggact
ggtcgtcggaacacgcattcggccttctggagaaataccagaatcaaatgctttgcttcaacgacgatatccaaggtacaggagccgtgatcctgt
ccggtgtcatcaacgcaattcggaaagtcgaaaaggagaatcaggtttcgcccagagaccaccgtattgtcttttacggagcaggctctgccgcc
atcggcgttgctcgtcagatccagtcctatttccagatcgaacacaacatgacggaggaagaggcaaagcacgtgttctggatcgtcgattctaaa
ggcctcgtcaccacaacacgtggtgataagctggctcagcataaagtctactacgctagaggcgataacgagggacaacagtataaagagctga
tcgacattgttaactataacctttatagcctcattggactcagcagcacgactggtgcatttaatacgcaggttctggaacgacttgcttcccttaatga
gcagcccattgtttttcctctgagcaatcctgcaacgcaggctgagtgtacctttgaacaggctatggaggctaccaataataaggttatctttgccag
cggaacagctttccccgcttatacgattaaatctacaggagaggtgaatacgcccggtcaaggcaataacatgtacatttttcccggccttggtctcg
gcgcatgccttgctaatcctgcacacttcgaccgtatgatctacgaagcctccaaagcactcgcagattccctgacagaagaagaaattagcaaag

cttggctctatccttctctcaactatcggagcgtttccgctatcgtcgccgccgcagtttgccaagagactctgaacgaaaatcttgccacctcccaa
gctatgatgacacaatgtaaatcgcatgaggatattcttgactatgttagcgctcacatgtggagccctgattacggaaataataattccaaccagca
agccggtaagtaa

**Appendix: LC-MS characterization of OLA standards and samples**

### Olivetolic Acid by LC-MS/MS

MCAC

For: Jingbo Ma (CBEE)
January 7, 2020

Report prepared by: M. LaCourse

Date: January 7, 2020

Report reviewed by: Joshua Wilhide

Date:

### Objective & Solution Preparation

- The goal of this analysis is to utilize LC-MS/MS to determine the olivetolic acid concentration in 3 samples
- Sample and standard preparation:
  - Samples were analyzed as provided by transferring to autosampler vials
    - Sample 3 was non-homogenous and thus vortexed thoroughly before analysis but still separated quickly
  - A standard curve was created in methanol from the provided 10 ppm stock following the scheme below:

| C1 (ppb) | V1 (uL) | C2 (ppb) | V2 (uL) | Diluent (uL) |
|---|---|---|---|---|
| 10,000 | 150 | 1000 | 1500 | 1350 |
| 10,000 | 50 | 500 | 1000 | 950 |
| 1,000 | 250 | 250 | 1000 | 750 |
| 1,000 | 100 | 100 | 1000 | 900 |
| 1,000 | 50 | 50 | 1000 | 950 |
| 1,000 | 10 | 10 | 1000 | 990 |

### Instrumentation & Method

**Instrumentation:**

- Qsight LX50:
    - PerkinElmer UHPLC Precision Sampling Module
    - Perkin Elmer UHPLC Solvent Delivery Module
    - PerkinElmer UHPLC Column Temperature Module
- Perkin Elmer QSight 210 Mass Spectrometer
- Column: Agilent Zorbax Eclipse XDB-C18 2.1 x 50 mm, 1.8 µm (P/N 981757-902)

**HPLC Method:** Olivetolic Acid V1

- Injection Volume: 10.0 µL
- Column Temperature: 40°C
- Run time: 5.00 min
- Flow: 0.3 mL/min
- Isocratic: (40:60 water:methanol)

**MS Method:** Olivetolic Acid V1

- Drying Gas: 80.0
- HSID Temperature (°C): 320.0
- Nebulizer Gas 1: 100.0
- ElectroSpray V1 Pos: -3000.0
- Source 1 Temperature (°C): 200.0

**Olivetolic Acid** (negative mode)

- Q1 Mass: 223.5
- Q2 Mass: 180
- Dwell Time: 795 ms
- Resolution (Q1:Q2): Unit_Unit
- CE: 22, EV:-30, CCL2: 20

### Calibration Curve

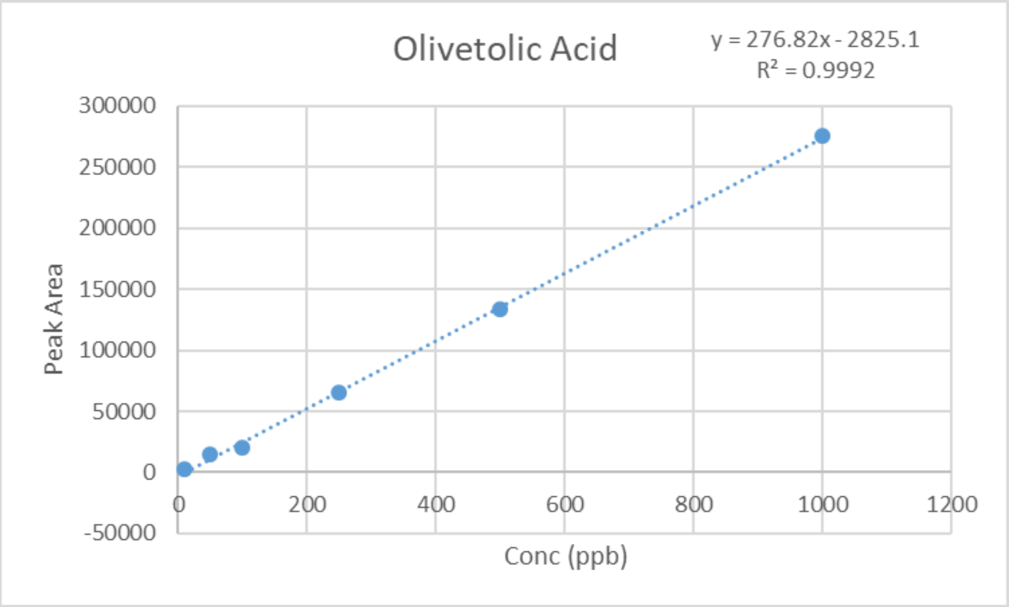

#### Methanol Blank

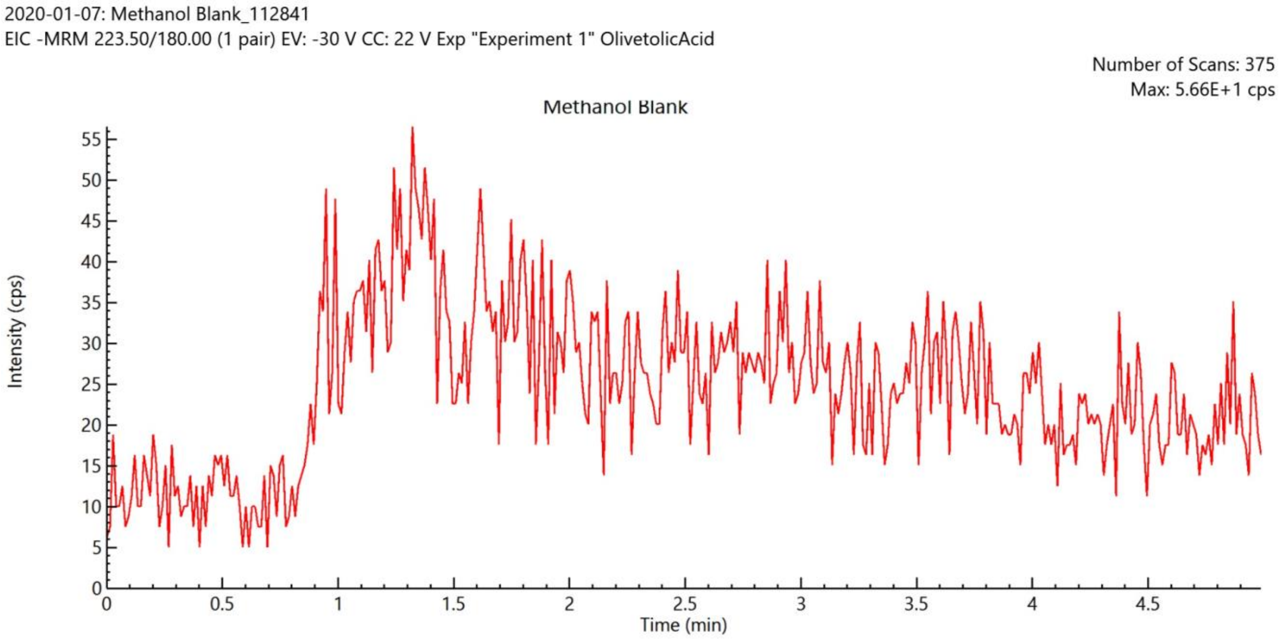

#### 500 ppb Olivetolic Acid Standard

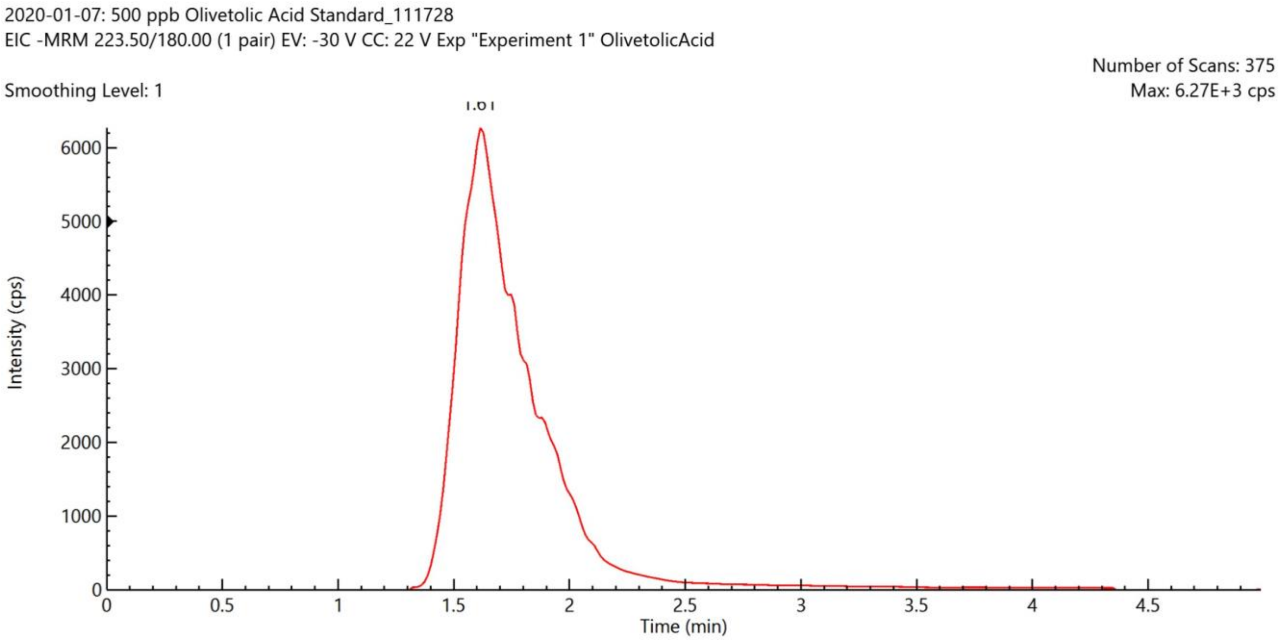

### Sample 1

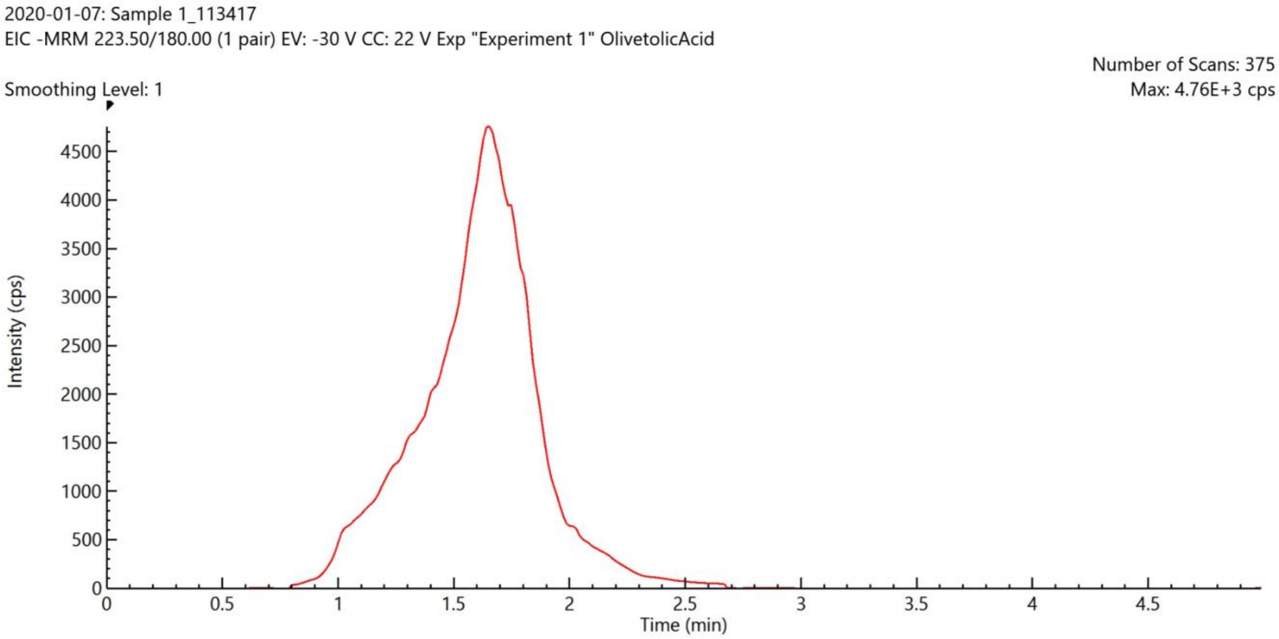

### Sample 2

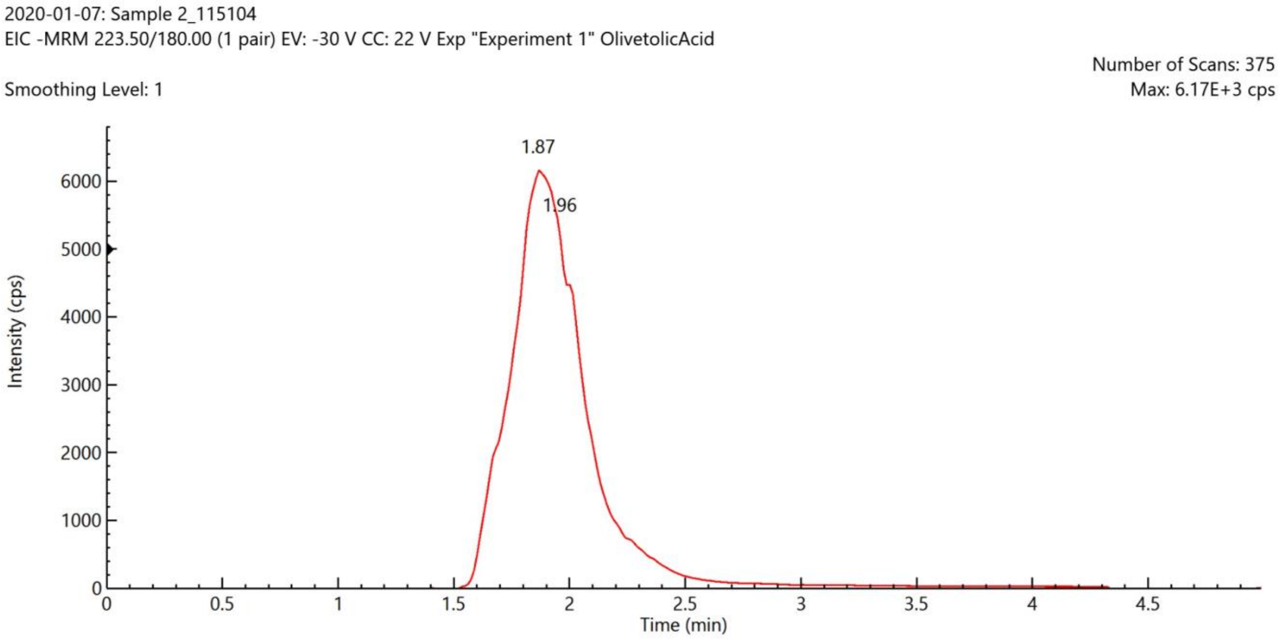

Sample 3

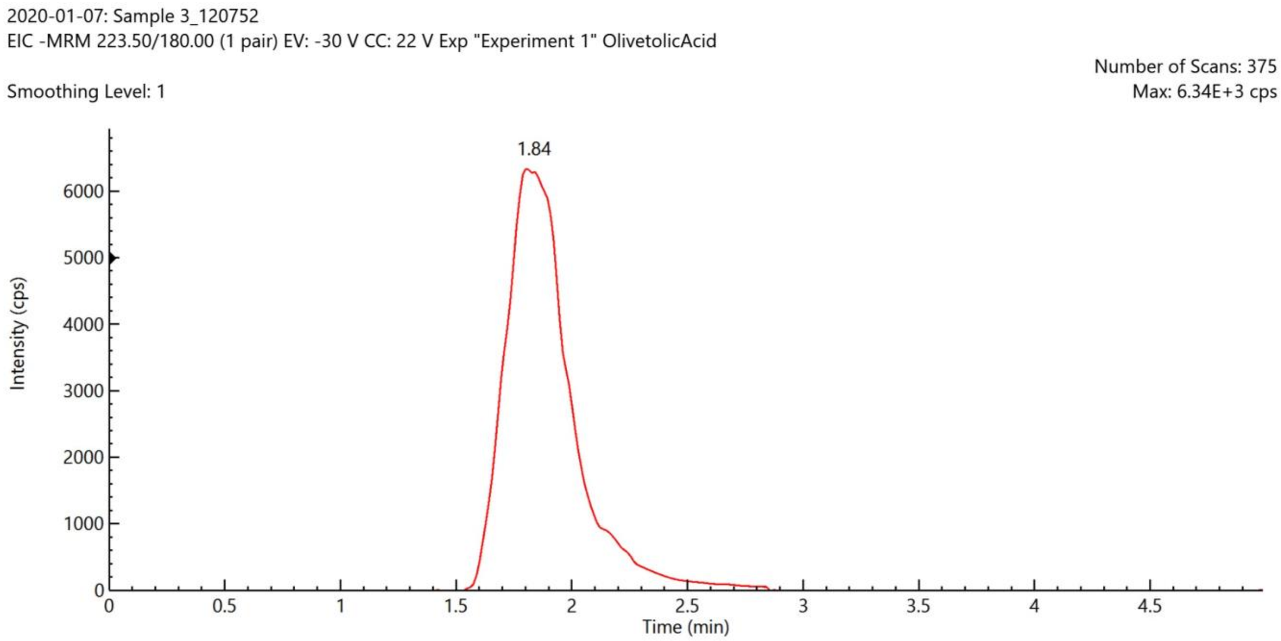

### Results

- See corresponding spreadsheet for all raw values and calculations

- Determined concentrations:
  - Sample 1: 649 ppb
  - Sample 2: 533 ppb
  - Sample 3: 467 ppb

### Olivetolic Acid by MS/MS

For: Jingbo Ma (CBEE)
January 14, 2020

Report prepared by: M. LaCourse

Date: January 14, 2020

Report reviewed by: Joshua Wilhide

Date:

### Instrumentation & Method

**Instrumentation:**
- Perkin Elmer QSight 210 Mass Spectrometer

**MS Method:**
- Drying Gas: 80.0
- HSID Temperature ($^{o}$C): 320.0
- Nebulizer Gas 1: 100.0
- ElectroSpray V1 Pos: -3000.0
- Source 1 Temperature ($^{o}$C): 0.0

**Olivetolic Acid** (negative mode)
- CE: 22
- Fragmentation:
  - EV:- 30
  - CCL2: 20

Mass Spectrum 1 ppm Olivetolic Acid

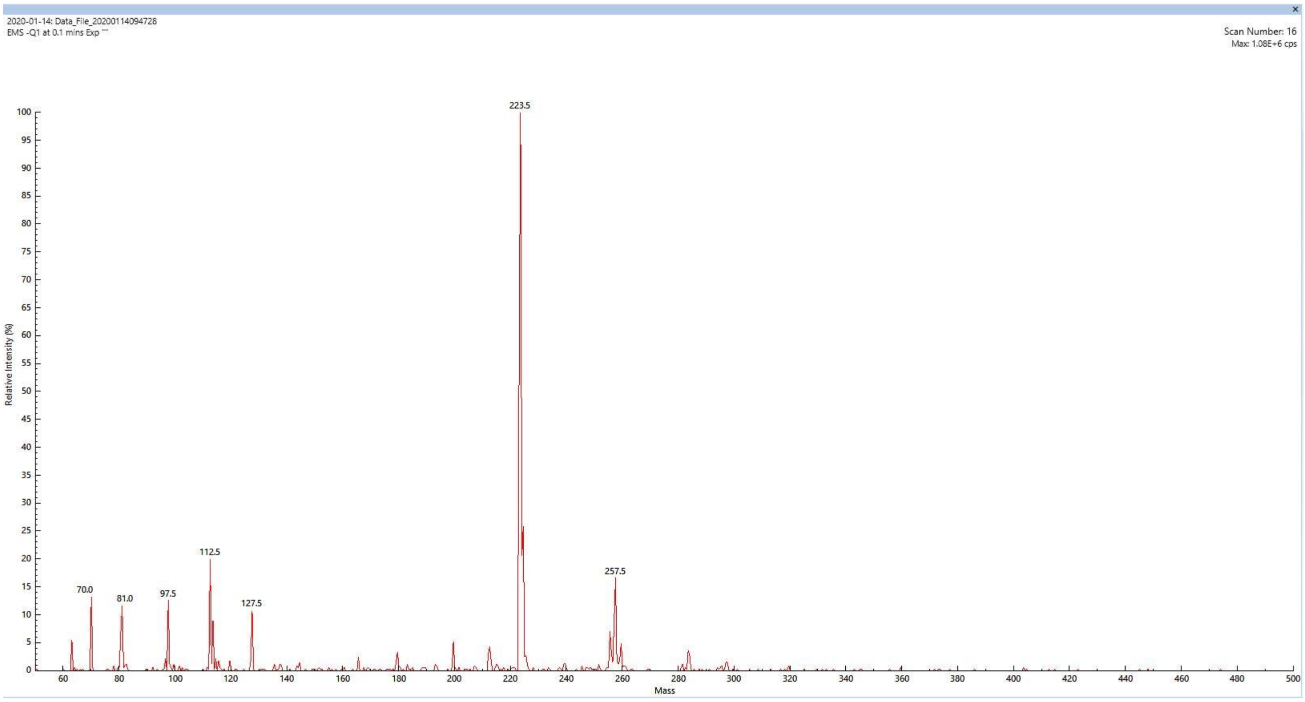

Mass Spectrum 1 ppm Olivetolic Acid - Fragmentation

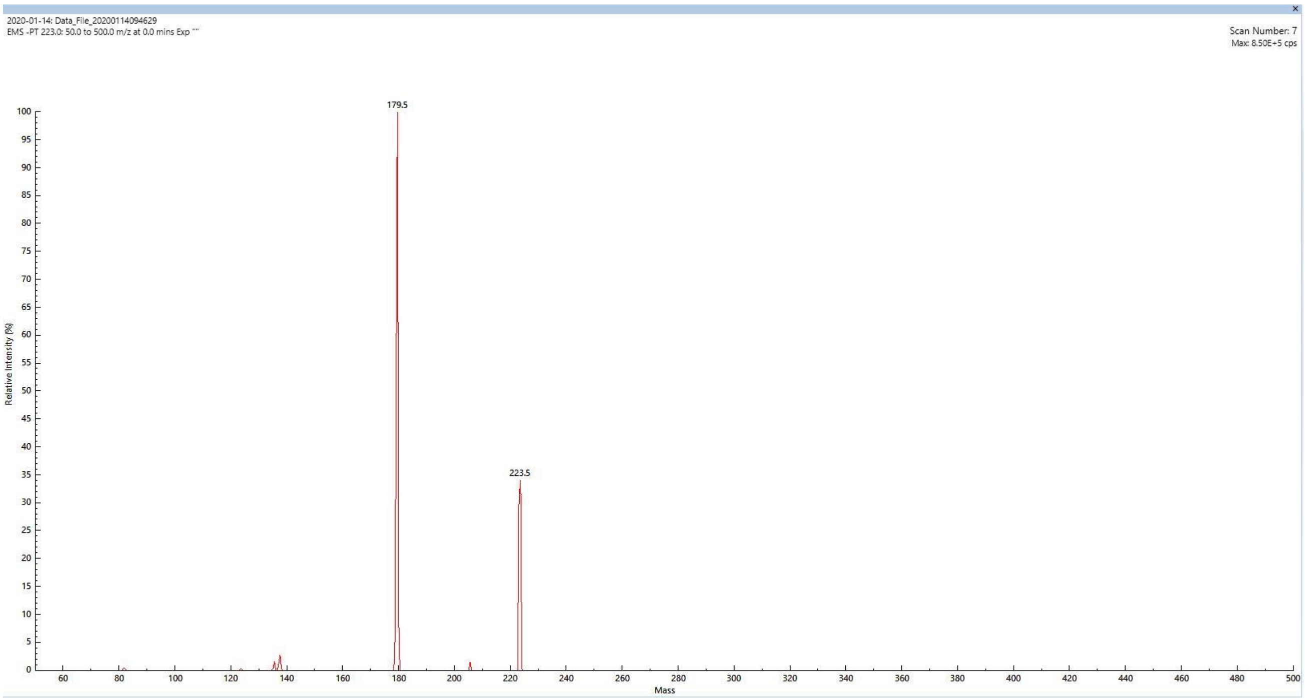

### Olivetolic Acid by High Resolution Mass Spectrometry (HRMS)

MCAC

For: Jingbo Ma (CBEE)
January 15, 2020

Report prepared by: M. LaCourse

Date: January 15, 2020

Report reviewed by: Joshua Wilhide

Date:

### Instrumentation & Method

**Instrumentation:**
- Bruker 12T solariX  FT-ICR-MS

**MS Method:**
- Negative mode ESI ionization
- Fragmentation with collision energy 12V

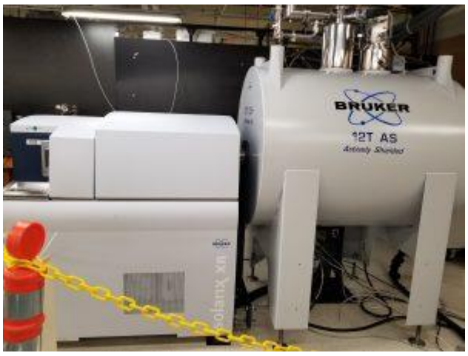

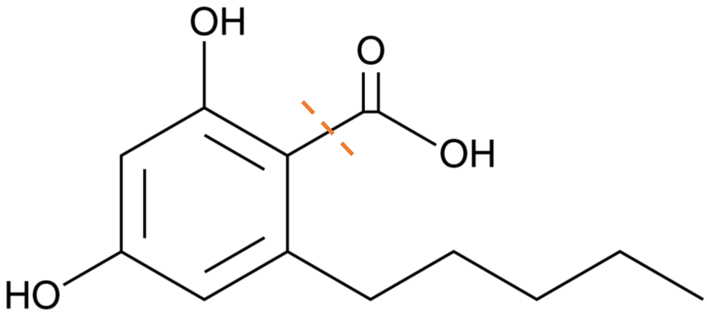

**Fragmentation**

$C_{12}H_{16}O_4 - COO \rightarrow C_{11}H_{16}O_2$

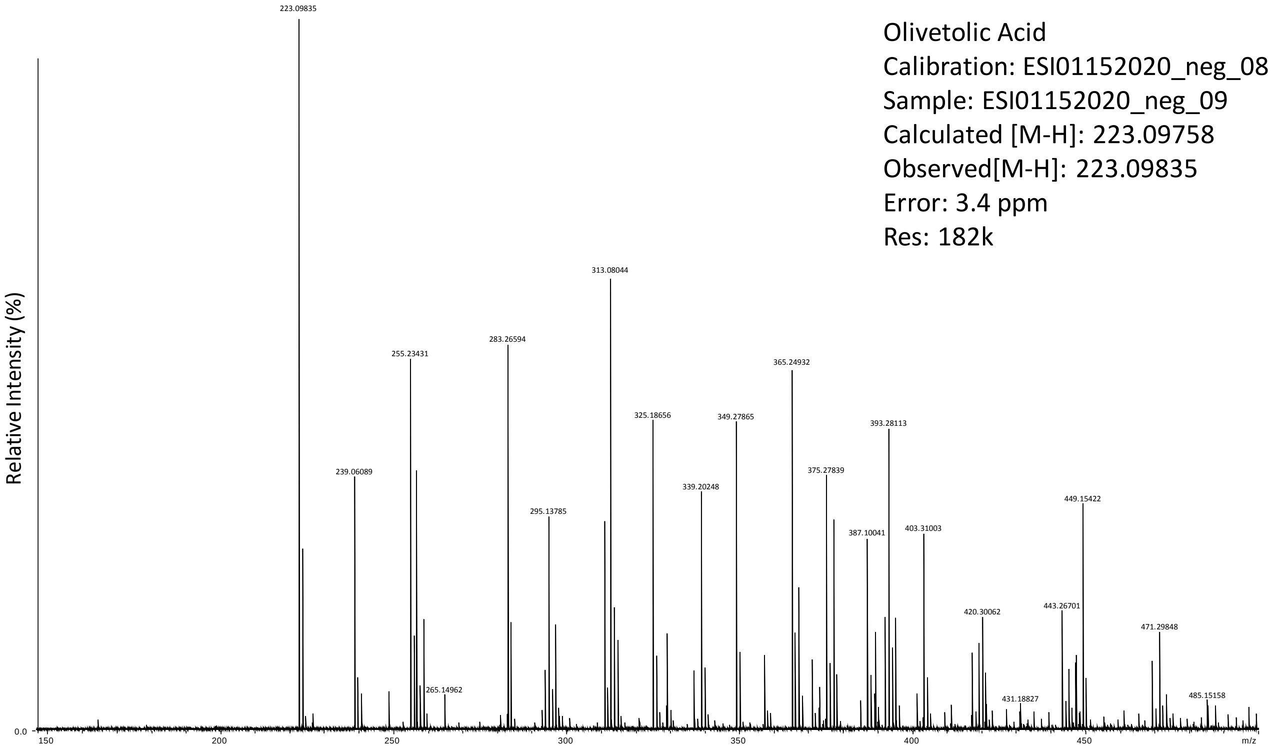

Olivetolic Acid
Calibration: ESI01152020_neg_08
Sample: ESI01152020_neg_09
Calculated [M-H]: 223.09758
Observed[M-H]: 223.09835
Error: 3.4 ppm
Res: 182k

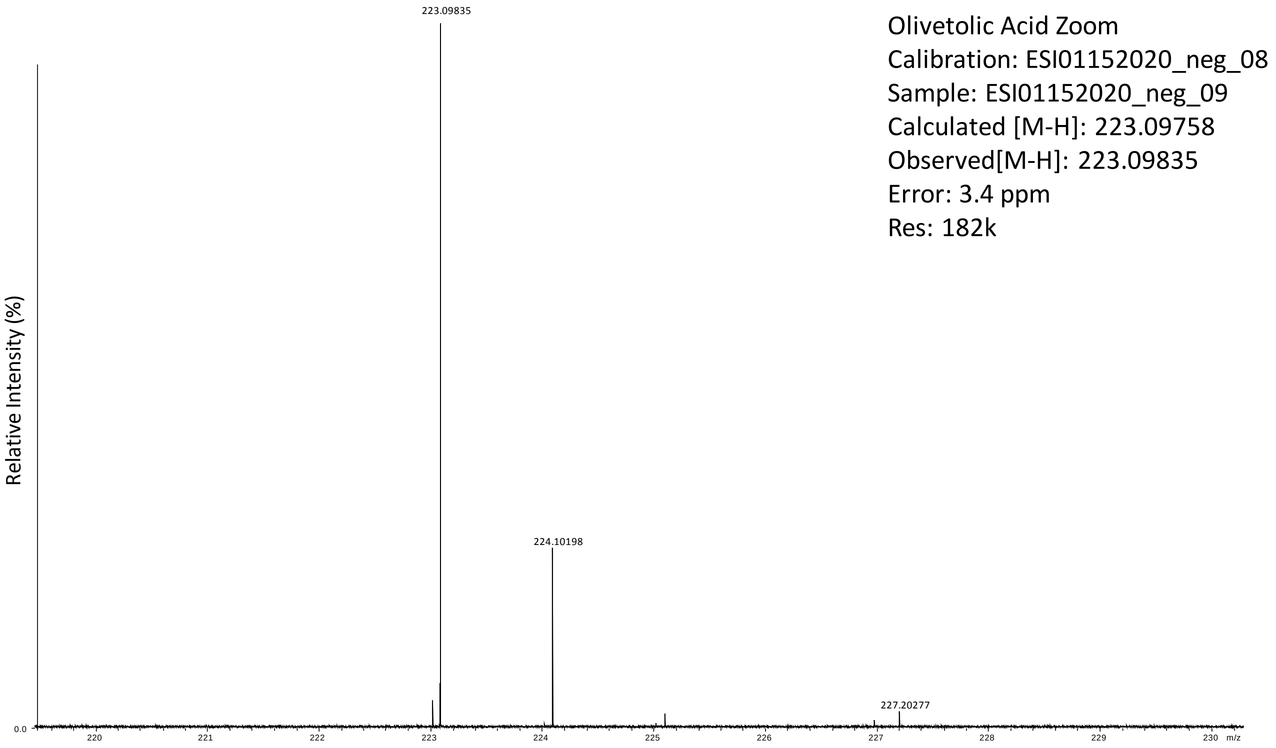
Olivetolic Acid Zoom
Calibration: ESI01152020_neg_08
Sample: ESI01152020_neg_09
Calculated [M-H]: 223.09758
Observed[M-H]: 223.09835
Error: 3.4 ppm
Res: 182k
Relative Intensity (%)
223.09835
224.10198
227.20277
0.0
m/z

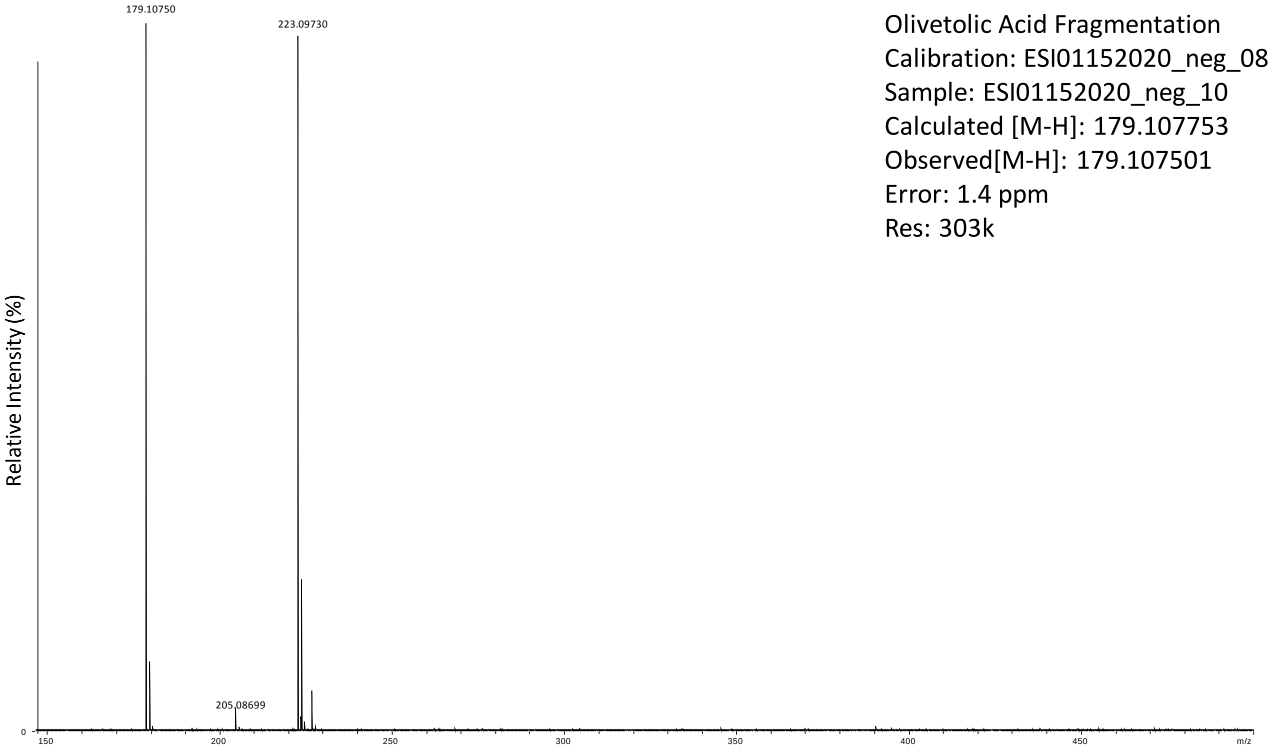
Olivetolic Acid Fragmentation
Calibration: ESI01152020_neg_08
Sample: ESI01152020_neg_10
Calculated [M-H]: 179.107753
Observed[M-H]: 179.107501
Error: 1.4 ppm
Res: 303k
Relative Intensity (%)
179.10750
223.09730
205.08699
0
150
200
250
300
350
400
450
m/z

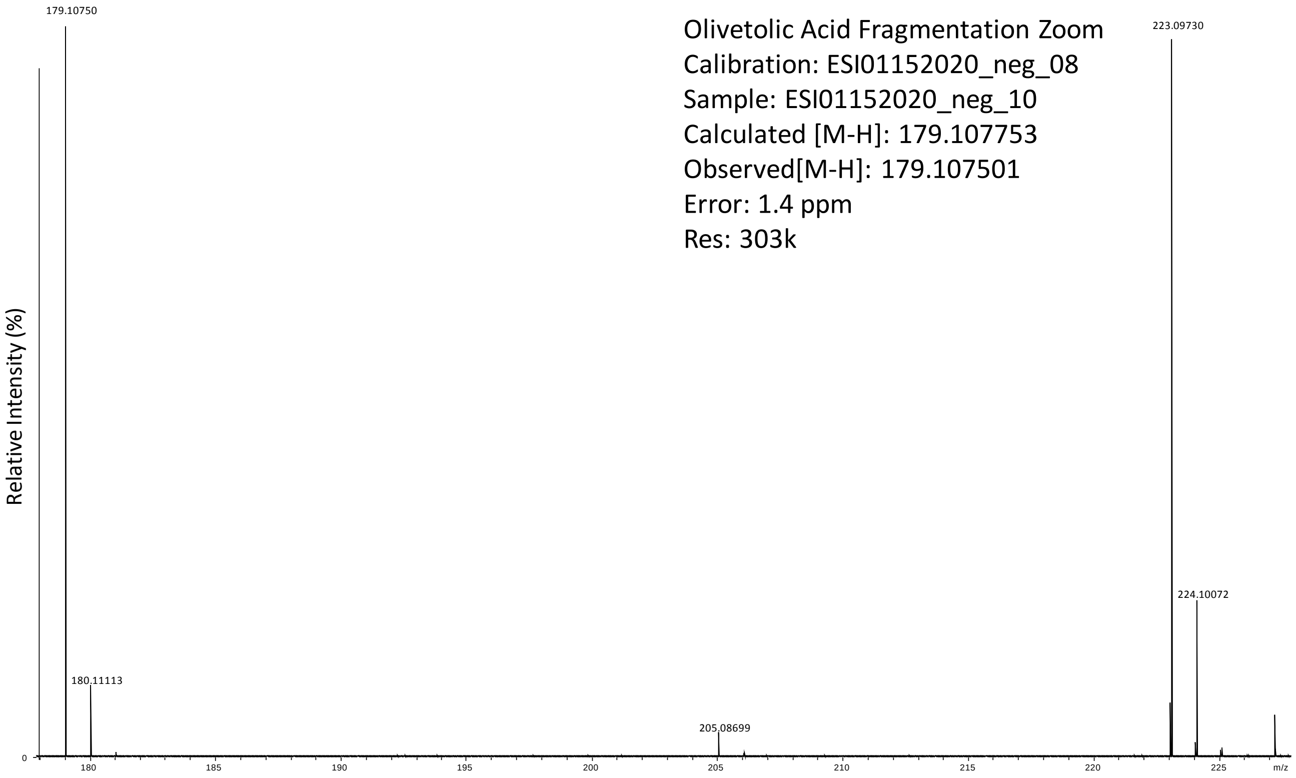

179.10750
Olivetolic Acid Fragmentation Zoom
Calibration: ESI01152020_neg_08
Sample: ESI01152020_neg_10
Calculated [M-H]: 179.107753
Observed[M-H]: 179.107501
Error: 1.4 ppm
Res: 303k
223.09730
224.10072
180.11113
205.08699
Relative Intensity (%)
0
180
185
190
195
200
205
210
215
220
225
m/z
